# Translocation of HIV capsid core through the Nuclear Pore Complex by affinity gradient

**DOI:** 10.1101/2025.10.02.680059

**Authors:** Ivo Melčák, Ryan L. Slack, Zachary C. Lorson, Andres Emanuelli Castaner, Krisztina Ambrus, Jonathan S. Winkjer, Karen A. Kirby, Robert A. Dick, Stefan G. Sarafianos

## Abstract

The HIV capsid core encapsulates the viral genome for subsequent integration into host cellular DNA. Prior to nuclear entry, the core must translocate through the Nuclear Pore Complex (NPC). This transit involves interactions between the capsid core and phenylalanine-glycine (FG) repeats found in nucleoporins within the NPC. Despite this critical role in the viral replication cycle, the molecular mechanism of capsid core translocation remains unclear. FG repeats consist of three classes of canonical motifs: FG, GLFG, and FxFG motifs. These are segregated within the NPC to define distinct zones of the gating machinery. FG- and FxFG-type motifs are enriched in the cytoplasmic and nuclear (“nuclear basket”) peripheries of the NPC while GLFG motifs are enriched in regions adjacent to the core of the NPC. To investigate the capsid core translocation, we use biochemical, biophysical, and structural approaches to study FG-capsid interactions in a quantitative manner. We show that the capsid (CA) interacts with a range of diverse FG repeats with varying affinities. GLFG motifs of core NUP98 exhibit increased affinity to CA proteins compared to other conventional FG/FxFG. However, the non-canonical FxFG motif of NUP153 at the “nuclear basket” significantly increases binding affinity to CA compared to canonical FxFG, therefore called FG super-motif. In addition, C-terminal motif of NUP153 consists of a stretch of basic residues, which enhances the affinity of this non-canonical FG super-motif to capsid core at the NPC nuclear periphery. We identified other binding enhancers of the NPC core FG-NUPs, NUP58 and POM121. The relationship between the binding strength of FG/FxFG binding enhancers of NUP58, POM121, NUP153 and their position within the NPC also shows capsid core binding affinity increases with increasing proximity to the “nuclear basket.” Based on our data, the difference in binding affinities between the canonical FxFG motif and the enhancer-coupled FG super-motif of NUP153 to capsid cores at the “nuclear basket” is approximately 1,000-fold. Therefore, the diverse FG motifs and binding enhancers, which are naturally distributed within the NPC into distinct zones, create an avidity gradient—with changes in both concentration and binding affinity—along the cytoplasmic-nuclear axis. We suggest that HIV capsid translocation into the nucleus is potentiated by this gradient in a unidirectional manner (outside-to-inside) within the NPC.

## INTRODUCTION

Human immunodeficiency virus type 1 (HIV-1) capsid core forms a compartment necessary for reverse transcription of its viral genomic RNA^1,2^. Following HIV fusion with the cell plasma membrane, the capsid core is released into the cytoplasm and interacts with capsid-targeting host factors that promote critical steps of entry into the nucleus where the viral cDNA product of reverse transcription is stably integrated into gene-rich regions of chromatin^3–7^. Recent evidence suggests that the capsid core can cross the NPC and subsequently disassembles in the nuclear interior^8–13^. Whereas HIV-1 reverse transcription and integration are relatively well understood, the mechanism by which a largely intact HIV capsid passes through the NPC remains to be elucidated.

The mammalian NPC is assembled from multiple copies of ∼30 nucleoporins (NUPs)^14^ and is anchored in the pore membrane domain of the nuclear envelope by distinct integral membrane proteins. The core of the NPC exhibits 2-fold symmetry in the plane of the nuclear envelope and 8-fold symmetry in the cytoplasmic-nuclear direction^15,16^. Attached to both sides of this symmetric core are filaments that are assembled into a “nuclear basket”^17^ on the nucleoplasmic side. The symmetric core houses the central transport channel with a reported diameter of ∼60 nm^8,18,19^.

Critical aspects of NPC function are collectively mediated by natively unstructured, phenylalanine-glycine (FG)-containing regions, which make up about a third of the NUPs^14,20^. Each FG region (150-700 amino acids in length) consists of multiple FG repeats separated by very hydrophilic, sometimes charged spacer sequences of 10-20 amino acids that lack a consensus sequence. These FG repeat domains are typically disordered, very flexible^21^ and come in different categories, distinguished by the specific nature of their phenylalanine-containing repeats and the composition and size of their spacers. FG repeats consist of three general classes of canonical motifs: 1) FG motifs, 2) GLFG motifs, and 3) FxFG motifs^22,23^. Extending from the scaffold to both the center and the sides—cytoplasmic and nucleoplasmic—of the central transport channel, FG repeats serve as a permeability barrier to restrict the passage of large molecules^20,24^ across the NPC and act as docking sites for the Karyopherin-β (Kap) family of nuclear transport receptors (NTRs)^25^. For a Kap to move its cargoes between the nucleus and cytoplasm in the correct direction, it must first bind the cargo in the starting compartment. The Kap-cargo complex then crosses the NPC by interacting with FG repeats. In the destination compartment, the cargo must be released from the Kap. The direction of Kap-mediated transport is regulated by RAN GTPase^26^. RAN-GTP and RAN-GDP are asymmetrically localized in the nucleus and RAN-GDP in the cytoplasm respectively. Kaps bind import cargoes and RAN-GTP in a mutually exclusive manner. Thus, Kaps bind import cargoes in the cytoplasm, translocate through the NPCs to the nucleus, where they bind RAN-GTP and release the cargo. The directionality of cellular cargo transport depends solely on the binding and unbinding influenced by the presence or absence of RAN-GTP on either side of the nuclear envelope^27–30^, rather than on an affinity gradient between Kaps and FGs.

Kaps mediate most cellular macromolecule transport, especially proteins, across the NPC^31–34^. Using X-ray protein crystallography, nuclear magnetic resonance spectroscopy (NMR) and molecular dynamics (MD) simulations, it has been shown that the bulk of FG-regions interact with NTRs predominantly through their FG motifs (phenylalanines within the FG motifs), and minimally through their spacer residues that separate FG repeats^35–40^.

The mature capsid core consists of capsomeres that comprise approximately 250 CA (capsid protein) hexamers and 12 CA pentamers, which assemble into a fullerene-like cone measuring about 120×50 nm^1,41^. Similar to cellular NTRs, HIV-1 capsid cores can directly bind FG repeats^42,43^ through a unique cavity called the FG-binding pocket. This pocket is a mainly hydrophobic cavity located at the NTD-CTD interface between adjacent monomers within a CA hexamer. It is primarily formed by hairpin residues of CA α helices 3 and 4. Structural studies have shown that NUP153 FG-containing peptide binds to the FG-binding pocket between two CA subunits of the hexamer^44–46^. Other host factors, including the primarily nuclear protein cleavage-and-polyadenylation-specificity factor 6 (CPSF6), also bind to the FG pocket of the core *via* the FG motif at the interface between two CA subunits^45–48^. These interactions influence HIV capsid nuclear transport^49–51^. It is proposed that CPSF6 competes with NUP153 for the common binding site on CA^44^, resulting in the capsid core being released into the nucleus^8,9,50^.

Recently, NUP153 was shown to interact with CA assemblies beyond hexamers at a much higher affinity. This strong interaction is achieved through a bipartite motif of NUP153, containing both the FG-motif and a C-terminal basic, triple-arginine (RRR) motif^52,53^. The RRR-motif binds at the interface of three CA hexamers, which is at the center of the 3-fold symmetry axis on the assembled capsid. This binding mode supports that the intact HIV capsid translocates through the NPC and disassembles in the nuclear interior^8–13^.

The high multiplicity of FG-binding pockets (∼1,500) and their interaction with FG repeats make the passage of the core particle through the NPC so forceful that it eventually damages the structural scaffold of the nuclear pore^9^.

Over the course of more than two decades, numerous models have been proposed to explain how disordered FG repeat regions form a diffusion barrier and cooperate to mediate nucleocytoplasmic transport. The structure of the permeability barrier — which partly depends on whether FG-NUPs interact with each other (are cohesive) or to what extent they repel each other (which could create brush-like structures) — still has not reached a consensus in the field (reviewed in^54^). The question of how the permeability barrier is constructed remains open and is beyond the scope of this study, which does not dependent on any of these models but can provide a conceptual framework for understanding the kinetics of HIV transport through the NPC. This framework relies solely on well-established features of nuclear transport receptor/FG-NUP binding that are common to all proposed models.

Mechanistic studies of HIV-1 nuclear entry are hindered mainly by the system’s inherent complexity. The NPC is one of the largest assemblies in eukaryotic cells (∼120 MDa^55^), and the HIV-1 capsid core is one of the largest cargoes (with a capsid core shell of ∼40 MDa) delivered to the cell nucleus^8–11^. While it is known that FG repeats of NUPs interact with the HIV-1 capsid^42,43^, the exact molecular mechanism by which the intact capsid translocates through the NPC remains unclear. Therefore, it is crucial to understand whether different FG sequences interact with capsid proteins in varying ways; here, we present detailed results on the physicochemical properties of HIV-1 CA interactions with various FG-containing moieties and bipartite FG motifs.

Based on the results of this study, we propose a model that explains the mechanism by which the HIV-1 capsid core is transported through the NPC. In our model, the gradient-based transport of the capsid core through the NPC relies on sequential interactions between FG repeats of NUPs and their binding enhancers, with affinities increasing gradually along the cytoplasmic-to-nuclear direction, from cytoplasmic filaments to the “nuclear basket”.

## RESULTS

Out of 10 major FG-NUPs, FG repeats fall into three main classes of canonical motifs: 1) FG motifs, 2) GLFG motifs, and 3) FxFG motifs^22,23^. These are organized within the NPC to establish distinct zones of the gating machinery. While FG- and FxFG-type motifs are concentrated at the cytoplasmic and nuclear (“nuclear basket”) peripheries of the NPC, GLFG motifs are more prevalent near regions adjacent to the central part of the NPC^56–58^ (**Figure 1**). Furthermore, in addition to the C-terminal basic RRR-motif^52,53^ located distal to C-terminal FG/FxFG repeats of NUP153 in the “nuclear basket” periphery, our primary structural analysis revealed a similar stretch of C-terminal basic residues near FG/FxFG repeats on two core FG-NUPs, NUP58 and POM121. This further emphasizes the functionally distinct zones within the NPC and contributes to the asymmetry of various FG-NUP “signatures” along the cytoplasmic-nuclear axis of the NPC.

**Figure 1.**
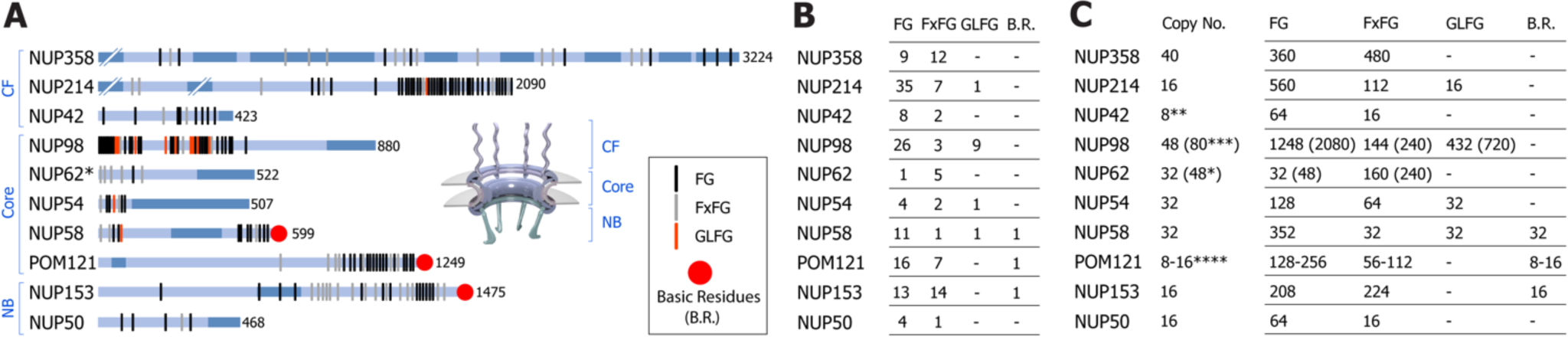
The catalogue of major human FG-NUPs within different zones of the NPC. (A) Structured domains and natively disordered regions are depicted in dark and light blue color respectively. C-terminal residue numbers are specified. Core NUP62 (*) is a promiscuous NUP as also a part of heterotrimeric cytoplasmic complex with NUP214 (and NUP88, not shown). Different FG flavors (vertical lines) and C-terminal basic residues (dots) are visualized and colored as indicated in the inset. Schematic representation of the structured NPC with cytoplasmic filaments (CF), core and the “nuclear basket” (NB) is shown in the middle. The structure of the NB is modified according to^17^. (B) Frequency of FG, FxFG, GLFG motifs and C-terminal stretches of basic residues (B.R.) for indicated FG-NUPs. (C) Stoichiometry (left) and total number of FG, FxFG, GLFG and motifs of C-terminal basic residues (B.R.) for designated FG-NUPs (right). 48 copies of NUP62 (*) include both cytoplasmic and core protomers. The consensus copy number of FG-NUPs per NPC is indicated, except 8 copies of NUP42 (**), 80 protomers of NUP98 (***), and 8-16 copies of POM121 (****) that are derived from^59,60–62^ respectively.

### Biochemical and biophysical characterization of CA-FG, CA-FxFG, and CA-GLFG interactions

The approximately 200 different human FG-repeats are separated by spacers that lack a consensus sequence^63^. To compare the binding strengths of FG, FxFG, and GLFG motifs on CA, we selected flanking sequences that are identical across all three motifs. We used natural FG-containing sequences from human NUP98, which are essential for maintaining the permeability barrier of the vertebrate NPC^64^ and contain all three motifs: FG, FxFG, and GLFG. We identified two native FG-containing repeats (NTG<u>GLFG</u>N and NTG<u>FSFG</u>N; NUP98_47–54_ and NUP98_297-304_ respectively) that share identical flanking regions but differ in FG types (GLFG *versus* FSFG, **Figure S1A**). Serine is most commonly found among all F*x*FG motifs in the human NUP proteome, and F*S*FG is a representative candidate in this study.

To prevent potential steric hindrance caused by the attached 6-FAM fluorophore, an additional sequence conservation analysis was performed, allowing addition of few variant amino acid residues to expand the flanking regions and the complete sequence of the competing peptides (**Figure S1B**). Consequently, the resulting 14-amino acid long peptides closely match a representative NUP98 natural sequence in terms of amino acid composition and conservation but feature different central FG, FSFG, and GLFG motifs.

Probing low-affinity interactions between proteins and ligands is challenging. Monodispersity and high solubility of the protein of choice to saturate the ligand are key factors for measuring their binding constants. Therefore, FG-, FSFG-, and GLFG-interaction studies with CA were performed using engineered, mutant CA proteins CA_A14C/E45C/W184A/M185A65_, which form monodisperse, highly soluble crosslinked hexamers in solution (≥ 1 mM). For simplicity, we refer to these hexamers as CA below. Binding affinities of CA hexamers with fluorescently conjugated FG-peptides (**Figure S2A**) were measured using microscale thermophoresis (MST). Consistent with reported data^39,66^, monovalent, single-motif interactions are relatively weak, with an apparent K_D_ around 1 mM, but are significantly stronger than a non-specific binding control (mutant FG peptide: FG to YG). However, differences in binding affinities of peptides containing FG, FSFG, and GLFG are clear (**Figure 2A**). The CA-GLFG interaction is the strongest (K_D_,_app_ ∼0.6 mM), followed by CA-FSFG (K_D_,_app_ ≥ 1.3 mM) and CA-FG (K_D_,_app_ ≥ 2.4 mM). Thus, the GLFG-derived peptide exhibits at least 2-fold and 4-fold higher affinity than FSFG- and FG-peptides respectively.

**Figure 2.**
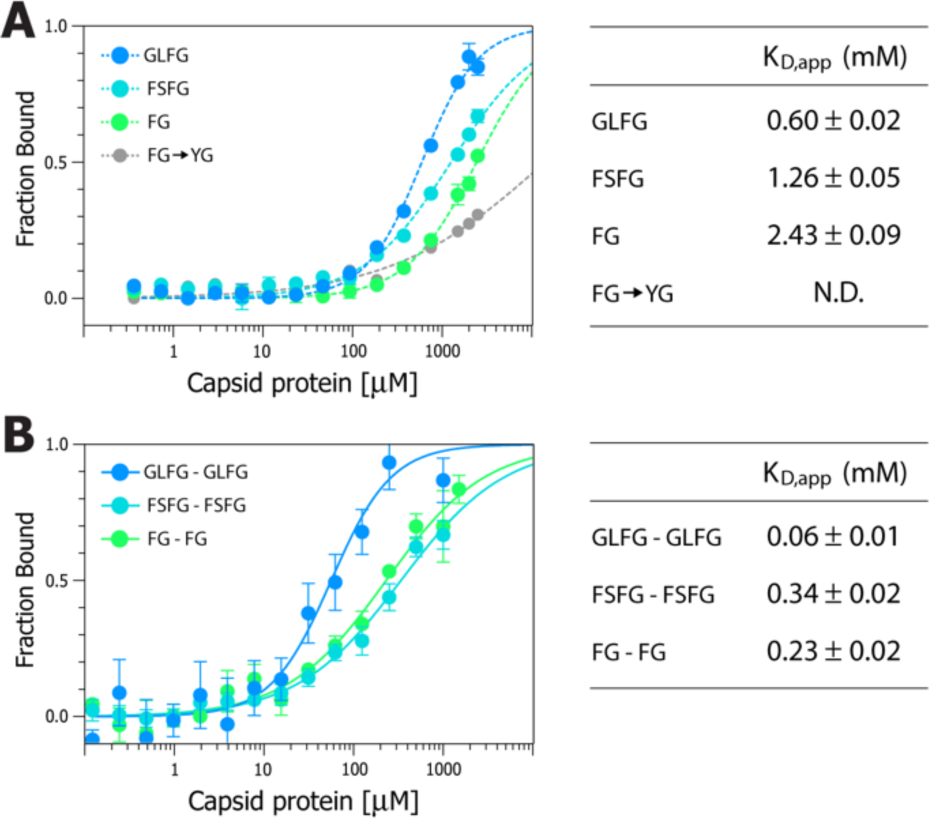
Affinities of different FG peptides to CA hexamers measured by MST. Binding profiles of peptides with single- (A) and double-motifs (B) of GLFG (blue), FSFG (cyan) and FG (green) are shown (left). Mutant FG to YG peptide is indicated in gray. Apparent K_D_ values (K_D_,_app_) are presented (right). N.D. not determined. Data are expressed as mean ± s.e.

When these motifs were dimerized, separated by a flexible ∼50-residue Gly-Ser repeat sequence (**Figure S2B**), interactions increased approximately 4 to 10 times compared to the corresponding monovalent peptides with the same tightest binder: CA-GLFG (K_D,app_ ∼0.06 mM) > CA-FG (K_D,app_ ≥ 0.23 mM) > CA-FSFG (K_D,app_ ≥ 0.34 mM) (**Figure 2B**). The lower avidity of the bivalent FSFG-derived peptide could be explained by a unique mode of interaction identified through structural analysis shown below. The enhancement of CA-GLFG binding compared to the other FG motifs (by roughly a further factor of 4) indicates that differences in binding affinities between GLFG and FG/FSFG are amplified through avidity.

In our MST assay, the measured affinities between CA and GLFG, FxFG, and FG do not necessarily reflect *in cellulo* values that would be observed in the crowded cellular environment. FG-repeats are exposed to multiple weak binders that significantly affect their interactions with binding parteners^67^. Accordingly, we performed MST experiments supplemented with *E. coli* lysate to examine the effect of such weak binders on the overall CA-GLFG avidity. As a prokaryote, *E. coli* does not contain nucleo-cytoplasmic transport proteins (e.g., Kaps) that would act as strong, natural competitors in our assay. In the presence of bacterial lysate, both monovalent and bivalent CA-GLFG interactions were attenuated (K_D_,_app_ N.D. and ∼0.2 mM respectively) but the cumulative strength of bivalent interactions was preserved (**Figure S3**).

Thus, even in a molecularly complex environment, GLFG-derived peptides bind CA with the highest affinity. Also, differences in binding affinities between GLFG and FG/FSFG are broadened through avidity.

### Using crystallography to determine the structural interactions of FG-, FSFG- and GLFG-peptides bound to CA

The goal of this study was to differentiate between FG, FSFG, or GLFG peptides with varying physicochemical properties bound to CA in an unbiased way. Therefore, identifying a single crystallization condition close to physiological conditions was ideal. We found one condition with pH and salt concentration near physiological levels that promoted the growth of co-crystals of crosslinked CA hexamers with either FG, FSFG, or GLFG peptides. Subsequently, we soaked these co-crystals with defined, higher concentrations of the specific peptides and determined their structures (**Figure S4** and **Table S1**). Since the relative affinities of monovalent “FG” peptides to CA are weak based on our MST studies, with K_D_,_app_ in the millimolar level, we utilized GLFG-, FSFG-, and FG-derived peptides with different, highly soluble flanking regions (see **Figure S2D**), as employed in other studies^68,69^. This allowed us to soak crystals with highly concentrated peptides, eventually saturating all binding sites in the crystal. Such approach was not possible with peptides containing “natural” flanking sequences employed in MST studies due to lower solubility.

Structural information from six crystal structures was obtained and used for semi-quantitative analysis of X-ray protein crystallography. Under the conditions used, the asymmetric unit of the crystals contains two CA hexamers, and each protomer has a unique FG-binding site. Therefore, for each crystal structure, twelve independent FG-binding sites were evaluated, which were not constrained by crystallographic symmetry. The binding profile of each peptide (defined by the visible density/structure) at a given apparent concentration was compared. In total, we obtained six structures of CA with various peptides containing diverse FG-motifs and concentrations, leading to the analysis of seventy-two FG-binding pockets (**Figures 3 and 4**).

**Figure 3.**
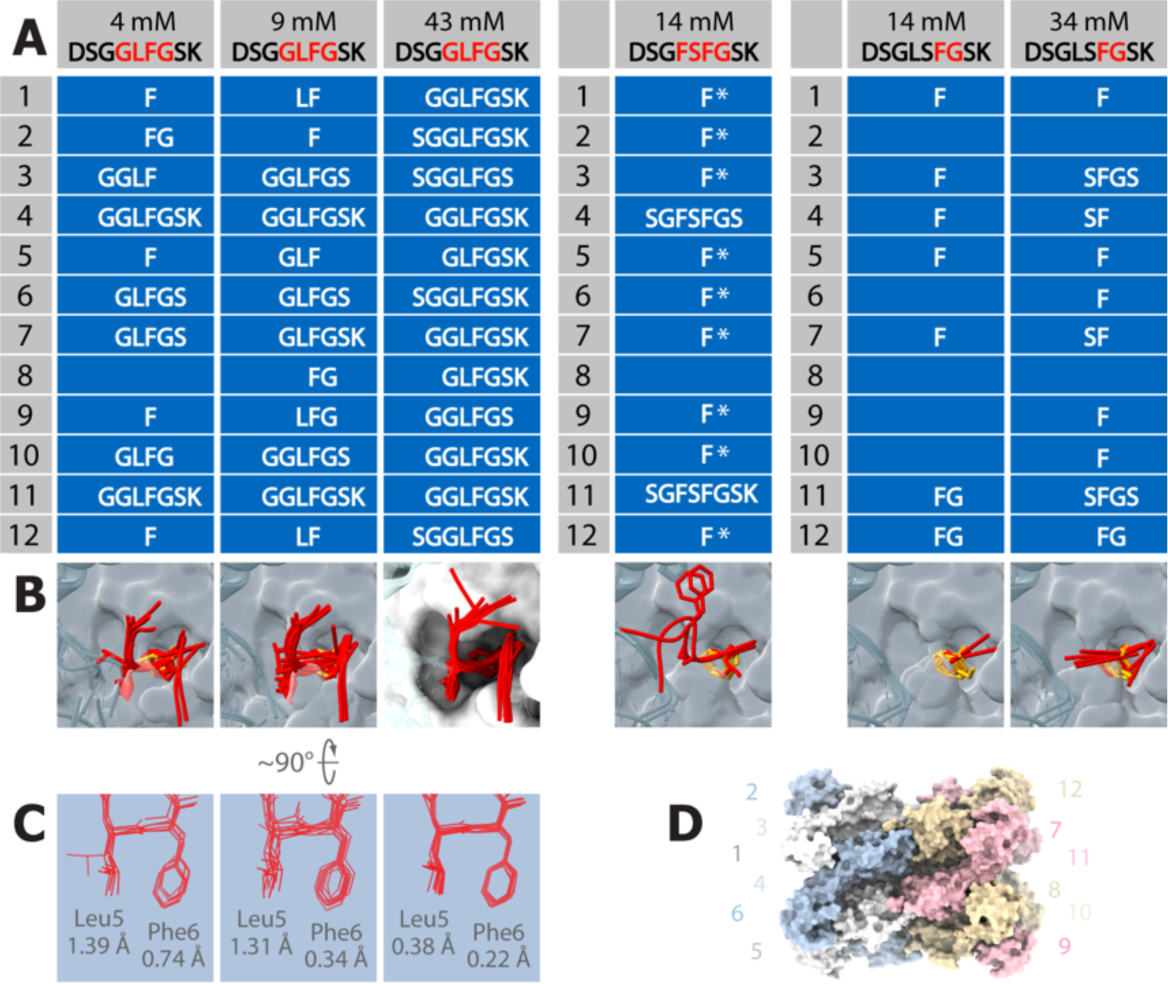
Structural interactions of FG-, FSFG- and GLFG-peptides bound to CA hexamer. (A) Table of peptide sequences derived from electron density maps for all twelve binding sites in the two CA hexamers of the asymmetric unit. GLFG-, FSFG- and FG-derived peptides (left, middle and right respectively). Apparent concentrations of peptides in crystal drops are indicated (top). The identity of which phenylalanine for FSFG is ambiguous (asterisk). (B) The superimposition of peptides (red) and sole phenylalanines (orange) from (A) onto binding pocket surfaces (transparent gray or solid white). Side chains for G<u>LF</u>G, <u>FSF</u>G and <u>F</u>G are shown in stick representation. The convergence of G<u>LF</u>G insertion from all peptides (43 mM) into FG-binding pockets (highlighted by overlayed solid white surfaces) is revealed. (C) Concentration-dependent RMSD values of overlayed leucine-phenylalanine containing structures (Leu5 and Phe6 from GLFG-derived peptides, shown in A). Values (in Å) decrease in concentration-dependent manner. (D) Schematic representation of crystal structures of CA hexamers that feature twelve unique FG-binding pockets. The asymmetric unit contains two CA hexamers in “head-to-head” orientation. Protomers are labeled with alternative colors (hexamer 1: blue-white, hexamer 2: pink-light brown).

**Figure 4.**
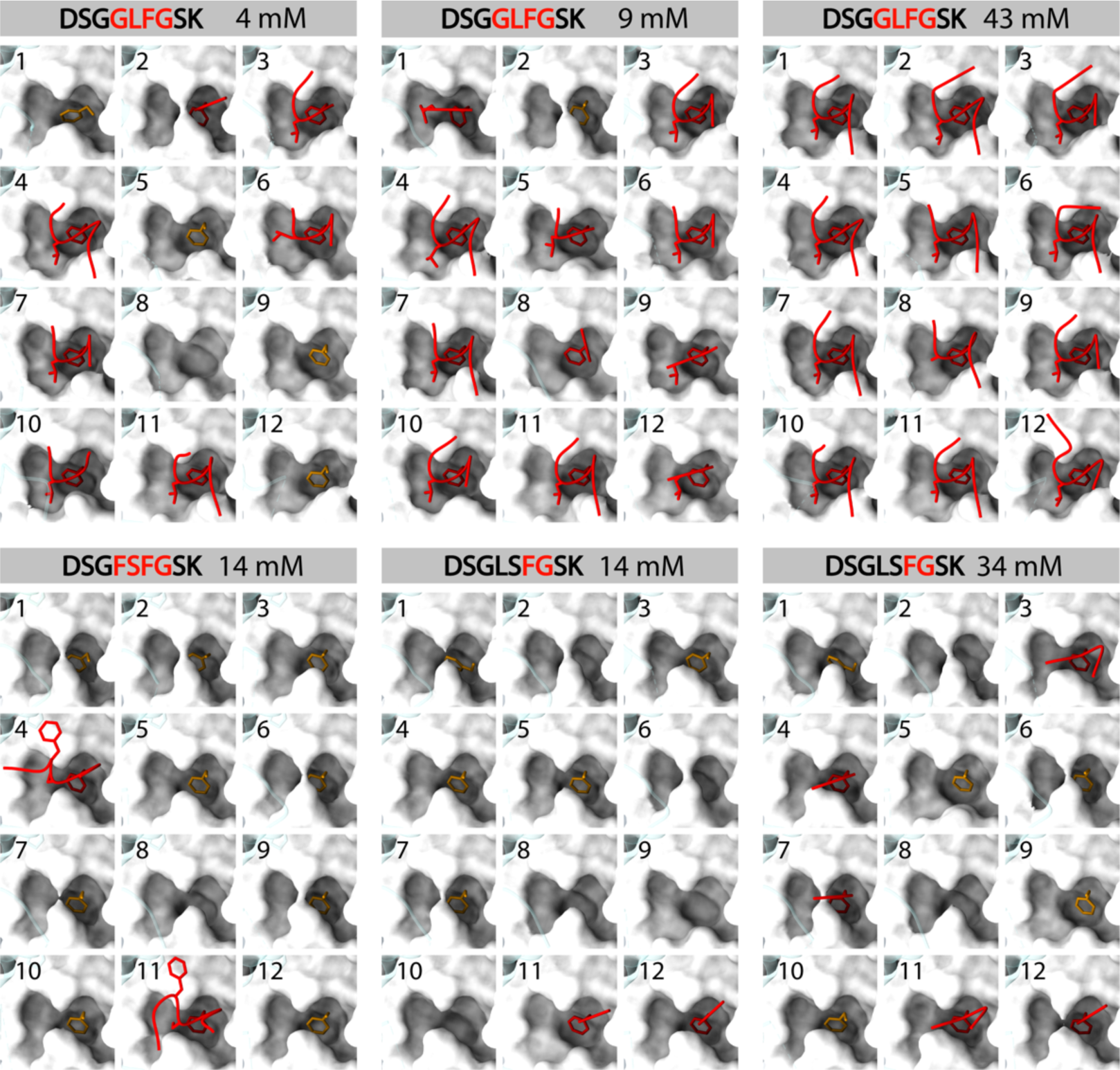
Structures of GLFG-, FSFG- and FG-peptides bound to CA hexamer. (Top) Occupation of CA FG-binding sites with GLFG-derived peptide (4, 9 and 43 mM; left, middle and right respectively). Twelve FG-binding pockets (1-12) of CA in surface representation with GLFG-containing peptides (red) and sole phenylalanines (orange). Side chains for G<u>LF</u>G are shown in stick representation. (Bottom) Occupation of CA FG-binding sites with FSFG-derived peptide (14 mM, left) and FG-derived peptide (14 and 34 mM; middle and right respectively). Twelve FG-binding pockets (1-12) of CA in surface representation with FSFG- and FG-containing peptides (red) and sole phenylalanines (orange). Side chains for <u>FSF</u>G and <u>F</u>G are shown in stick representation. Peptide sequences and apparent concentrations of peptides in crystal drops are indicated.

In general, we observed both quantitative and qualitative differences in peptide binding depending on the concentration, which enabled us to compare the relative affinities among FG, FSFG, and GLFG. First, FG-binding site occupation was concentration-dependent for all peptides studied (**Figures 3 and 4**). Second, the peptides fold into a stable CA-binding conformation in a concentration-dependent manner (**Figures 3A and 3B**). For example, the GLFG-derived peptide became progressively less disordered as the peptide concentration increased (4 mM, 9 mM, and 43 mM respectively). Presumably, the folding of GLFG- and FG-derived peptides occurs after the initial insertion of phenylalanine into the binding pocket. This idea is supported by the characterization of folded intermediates for these two motifs (**Figure 4**). In contrast, a single mode of interaction without visible intermediates was observed for CA-FSFG (**Figure 4**), likely reflecting differences in the kinetics of binding. This supports the MST binding studies described above, which show lower avidity of bivalent FSFG-peptides compared to GLFG- and FG-peptides. Third, the formation of stable secondary structure was accompanied by rigidification of FG-motifs in the binding pocket, and *vice versa*, as indicated by the consistently lower root-mean-square deviation (RMSD) values (**Figure 3C**). Based on these criteria, the relative monovalent affinities of FG-containing peptides were determined to be GLFG > FSFG > FG, in agreement with the MST data. As a complementary approach to confirm the occupancy of GLFG, FSFG, and FG in the crystal structures, we analyzed the position of Gln179 relative to the FG-binding pocket (**Figure 5**). Gln179 is situated in the loop between α helices 8 and 9 of the CTD of CA, adjacent to the NTD FG-binding pocket (α helices 3 and 4). We observed that the occupancy of the FG pocket by all three peptides and the position of Gln179 are correlated in a concentration-dependent manner (**Figure 5A**). In the unoccupied FG-binding cavity, the loop between α helices 8 and 9 is closer to α helices 3 and 4, and Gln179 readily occludes the pocket (**Figures 5B and 5C**). When peptides are bound, Gln179 is displaced from the binding pocket due to loop disorder, often accompanied by missing side chains or entire backbone density (**Figures 5D and 5E**). Only a small portion of Gln179 appears mislocalized because of crystal packing effects (**Figure 5F**). Based on this complementary approach, the measured displacement of Gln179 relative to the FG-binding pocket is consistent with a decreasing order of affinities: GLFG > FSFG > FG.

**Figure 5.**
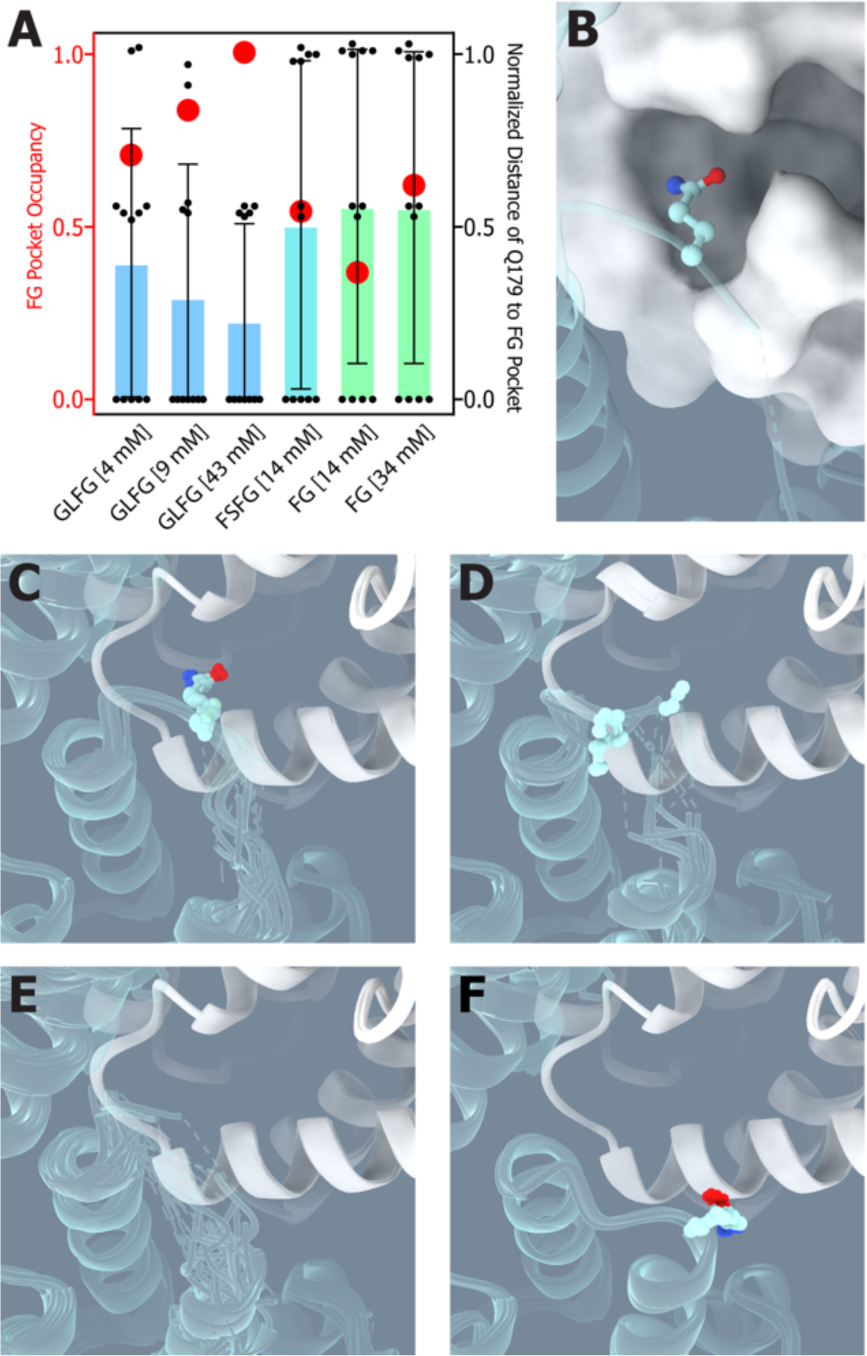
A correlation between GLFG, FSFG- and FG-peptide concentrations and occupancy of Gln179 in FG-binding pocket. (A) Comparison of FG pocket occupancy and distance between Gln179 and the FG binding pocket. Occupancy (left axis, red circles) is defined as the sum of Phe and Gly residues resolved in each independent crystal structure divided by the total number of Phe and Gly residues in the crystallographic asymmetric unit (24). Note that these values positively correlate with the peptide concentration used for crystallization. Distances between the C_α_ atom of Gln179 and the C_α_ atom of Asn57 within the FG binding pocket were calculated for each CA subunit and normalized with respect to this distance of Gln179 occluding FG-binding pocket (right axis, each distance is depicted as a black dot, colored bars represent the median of all 12 distances within the structure and error bars represent the standard deviation). In the case that C_α_ atom of Gln179 was unresolved, a normalized distance value of 0 was assigned. Note that the intra-subunit Gln179_Cα_-Asn57_Cα_ distance is negatively correlated with the peptide concentration used for crystallization, and FG pocket occupancy. (B) Representative figure of Gln179 (in ball-and-stick) in unoccupied FG-binding cavity (white surface). (C) Overlay of Gln179 obstructing FG pocket from all six X-ray crystal structures. (D) Same as in (C), but side chains of Gln179 are unresolved (represented by C_α_-C_β_ atoms only), and the loop is deflected from the binding pocket. (E) Same as in (C), but the whole loop between α helices 8 and 9 is disordered. (F) Overlay of mis-localized Gln179 (in ball- and-stick representation) from X-ray structures as a result of crystal packing.

We also found that the binding “footprints” of GLFG, FSFG, and FG motifs differ. Within the CA binding pocket, the flexibility of the Lys70 side chain allows the pocket to remodel itself to accommodate different motifs (**Figure S5**). Additionally, the larger binding surface of the GLFG motif compared to FG is mainly localized on one CA protomer, while the FSFG motif primarily connects two protomers through a stacking interaction between the first phenylalanine of <u>F</u>SFG and Pro38 of the adjacent protomer (**Figure S6**).

### Structural characterization of canonical and non-conventional FxFG-containing motifs bound to CA

The traditional (canonical) short FSFG motif used in our study shows low affinity for CA (K_D_,_app_ ≥ 1.3 mM). In contrast, the non-traditional, FTFG-derived motif from the C-terminus of NUP153 (NUP153_1411-1418_)^44,45^ exhibits increased binding compared to other NUP153 FG/FxFG-containing repeats^44,52^. The unique PSGV<u>FTFG</u> motif forms close contact between two protomers and extends the interaction beyond the FG-binding pocket through FTFG preceding residues, PSGV^45^. Comparing the CA-bound FSFG-derived peptide (**Figure 6**) with the PSGV<u>FTFG</u>-containing peptide from NUP153 (NUP153_1410-1418_)^45^ highlights the importance of specific flanking residues for binding. For canonical FxFG motifs, the most common first preceding amino acid is a small, flexible residue such as Gly, Ala, or Ser in over 60% of the major FG NUPs proteome (>60%). Gly is the most frequent (over 30%) and is used in our study (<u>G</u>FSFG), forming a turn outside the pocket. Conversely, the residue before FTFG in NUP153, Val1414 (*V*<u>FTFG</u>), helps bind the non-canonical PSG*V*<u>FTFG</u> motif by inserting into an additional pocket on the neighboring protomer (**Figure 6**). The mutation P1411A reduces binding to CA by about 10-fold (K_D_ ∼0.58 mM)^45^, consistent with our MST data showing weak, canonical FSFG-CA interactions, with an apparent K_D_ around 1 mM. Hence, we call this unique NUP153 sequence a non-canonical FG (or PSGV<u>FTFG</u>) super-motif.

**Figure 6.**
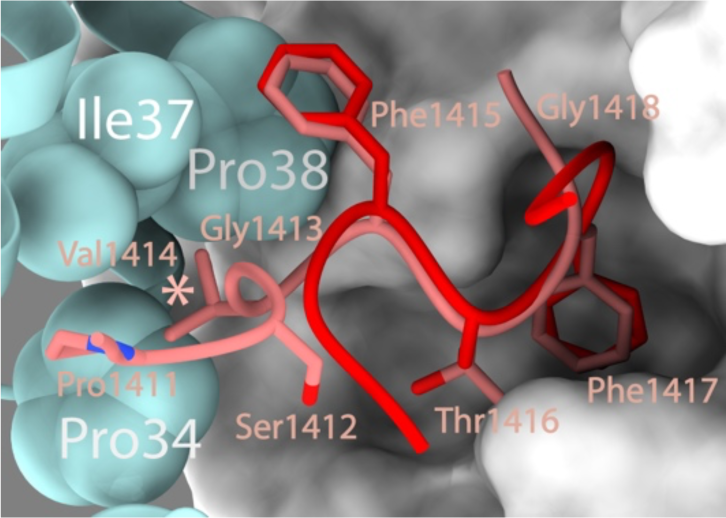
Canonical FSFG motif and PSGV<u>FTFG</u> super-motif in FG-binding pocket. Structural overlay of FSFG-derived peptide from our study (red) and non-canonical, FTFG-containing peptide of NUP153^20^ (NUP153_1410–1418_; pink) in the binding pocket. Phenylalanines are shown in stick representation. Val1414 of NUP153 (in stick, asterisk) binds to the pocket created by Pro34, Ile37 and Pro38 (spheres, labeled) of the adjacent protomer. The residues of PSGVFTFG are labeled.

### Characterization of interactions between FG-containing binding enhancers to CA assemblies

This unique, non-canonical PSGVFTFG super-motif from the C-terminus of NUP153 (NUP153_1411-1418_) exhibits stronger binding to CA than other nucleoporins’ FG/FxFG repeats^44,45,52^. This enhancing property is supported by a C-terminal triple-arginine (RRR) motif that interacts at the interface of three CA hexamers within a pocket of negatively charged residues^52^. Our primary structure analysis identified a stretch of basic residues near the C-terminal FG/FxFG repeats on two other FG-NUPs, NUP58, and POM121 (**Figure 7A**).

**Figure 7.**
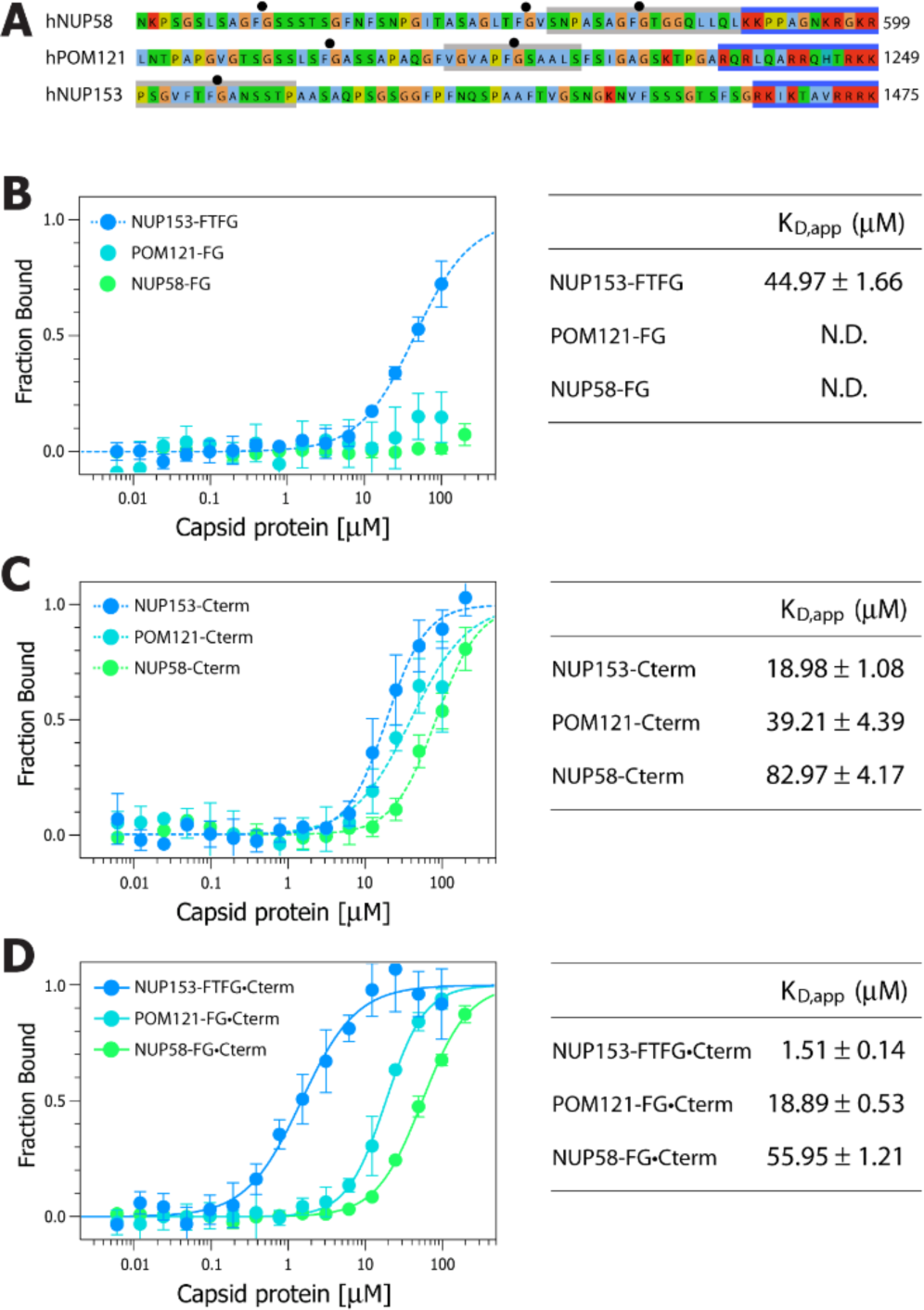
Characterization of enhancer-coupled FG/FxFG interactions. (A) Sequence analysis and design of FG-peptides with binding enhancers. Primary sequences of C-terminal regions of human NUP58, POM121 and NUP153. Basic residues are highlighted in red and FGs are indicated (black dots). Sequences underlaid with boxes represent those in engineered peptides (FTFG/FG in gray and basic-residues region in blue). Peptides used for MST assay and their designation, NUP58-FG•Cterm: NUP58_571-587/588-599_, NUP58-FG: NUP58_571-587_, NUP58-Cterm: NUP58_586-599_, POM121-FG•Cterm: POM121_1212-1223/1236-1249_, POM121-FG: POM121_1212-1223_, POM121-Cterm: POM121_1234-1249_, NUP153-FTFG•Cterm: NUP153_1411-1424/1465-1475_, NUP153-FTFG: NUP153_1411-1424_ and NUP153-Cterm: NUP153_1463-1475_. (B-D) Affinities of FG/FxFG motifs (B), C-terminal enhancers (C) and enhancers-coupled FG/FxFG (D) to CLPs. Binding profiles are shown (left). Color-coding is indicated and for designation, see (A). Apparent K_D_ values (K_D_,_app_) are presented (right). N.D. not determined. Data are expressed as mean ± s.e.

We tested whether these motifs also function as FG-binding enhancers using MST. Since the interface between three CA hexamers forms part of a higher-order lattice, we performed MST experiments with core-like particles (CLPs) that closely resemble authentic CA cores^70^ (**Figure S7**). First, we used fluorescently conjugated single FG-peptides without their C-terminal basic residues to measure their binding strength. These peptides include FG-containing NUP58_571-587_, POM121_1212-1223_, and non-canonical PSGVFTFG-containing NUP153_1411-1424_ (**Figure S8**). Consistent with our data (**Figure 2A**), naturally occurring FG-peptides of NUP58 and POM121 show low affinity, below the detection threshold of our MST assay. However, the non-canonical PSGVFTFG-containing NUP153 peptide exhibited much higher affinity, comparable with published data^45^ using crosslinked CA hexamers (K_D_,_app_ ≥ 45 μM and 49 μM; **Figure 7B**). Next, we tested the binding capacity of fluorescently conjugated peptides with C-terminal basic residues: NUP58_586-599_, POM121_1234-1249,_ and NUP153_1463-1475_, which contain the embedded RRR motif^52^ (**Figure 7C**). All three peptides show significant affinity to CLPs, with the increasing affinity order: NUP58_586-599_ (K_D_,_app_ ≥ 83 μM), POM121_1234-1249_ (K_D_,_app_ ≥ 39 μM), and NUP153_1463-1475_ (K_D_,_app_ around 19 μM). The somewhat lower binding strength of NUP58 may be due to the intervening glycine residue (KR<u>G</u>KR), which splits the C-terminal basic region.

Our data show that a non-canonical PSGVFTFG super-motif of NUP153 increases binding capacity to CA by approximately 30-fold compared to canonical FSFG (see **Figure 6** for structural comparison). Accordingly, the binding affinity is also enhanced by about 30-fold (**Figure 7D**) when this NUP153 peptide is extended with a C-terminal RRR motif (NUP153_1465–1475_). The resulting affinity (K_D_,_app_ ∼1.5 μM) is comparable to the reported value (K_D_ ∼0.9 μM)^52^ when assembly mimicking the interface between three CA hexamers was used. Increased affinity was also observed when FG-containing peptides of POM121 and NUP58 were lengthened with their C-terminal basic residues (POM121_1236-1249_ and NUP58_588-599_ respectively; see **Figure S8**). The corresponding K_D_,_app_ values for POM121 and NUP58 bipartite peptides were ≥ 19 μM and ≥ 56 μM (**Figure 7D**). The lower affinity of the NUP58 bipartite peptide compared to POM121 likely results from the weaker affinity of its C-terminal basic stretch to CLPs (K_D_,_app_ ≥ 83 μM and ≥ 39 μM respectively). Therefore, the positive correlation between the differences in affinities of individual monovalent and bivalent interactions indicates that overall binding strength is probably influenced by the “forced proximity”^71^ of FG/FxFG motifs to C-terminal basic residues.

In conclusion, the C-terminal basic residues of NUP58 and POM121 act similarly to those in NUP153. Thus, we refer to them collectively as (non-canonical) binding enhancers.

## DISCUSSION

HIV CA cores are specific nuclear transport receptors (NTRs) that show strict directionality^12,13^, moving from the cytoplasm to the nucleus. In contrast, cellular NTRs appear to shuttle back and forth through the NPC without directional bias^72^. The directionality of NTR-cargo import depends entirely on the chemical gradient of Ran-GTP which ratchets the cargo into the nucleus^27,73^. Although it is established that the phenylalanine-glycine (FG) repeats of a subset of NPC components, nucleoporins (FG-NUPs), interact with HIV CA^42,43^, the molecular mechanism by which the intact capsid translocates through the NPC remains not fully understood.

Based on the known structure and overall stoichiometry of FG-NUPs within the human NPC^18,60,74–78^, there are ∼5,000 various FG-repeats from multiple copies of ten major FG-NUPs (**Figure 1**). These FG-NUPs contain three main classes of canonical FG repeats: FG, FxFG, and GLFG. These motifs are localized within distinct zones of the NPC. The FG/FxFGs of NUP358, NUP214, and NUP42 protrude into the cytoplasm as part of the cytoplasmic filaments^76,79,80^. Attached to the core of the NPC^18,60,75–78^ are the FG/FxFGs of NUP58, NUP54, NUP62, POM121, and the GLFGs of NUP98. While most NUPs are present in 8-32 copies per NPC^14^, the structure of the human NPC core suggests 48 and possibly up to 80 copies of NUP98^60^. Therefore, GLFG nucleoporin NUP98 is one of the most abundant proteins of the NPC.

Our study shows that the GLFG motif of NUP98 exhibits increased affinity to HIV capsid proteins compared to FG/FxFG, and this difference in binding affinities between GLFG and FG/FxFG is compounded through avidity. In addition, we identified C-terminal binding enhancers on NUP58 and POM121. These FG/FxFG binding enhancers are present on 32 protomers of NUP58^18,60,75,77,78^ and 8-16 protomers of POM121^61,62^. Such opportunistic motifs adopted by the virus that bind capsid core directly further increase the binding potential of NPC core for HIV capsid.

On the nuclear side, the FG/FxFGs of NUP153 and NUP50 project to the nuclear basket^17,81,82^. Structural comparisons between the CA-bound, canonical, FxFG-containing motif and the reported PSGVFTFG-comprising peptide of NUP153^45^ reinforce the idea of HIV-specific binding motifs. Therefore, we refer to this unique PSGVFTFG sequence as a non-canonical FG super-motif. Our data indicate that this virus-sensing, non-canonical NUP153 FG super-motif increases binding capacity to CA by approximately 30-fold compared to canonical FxFG. This strengthening property is further enhanced by approximately 30-60 fold by the distal C-terminal binding enhancer, originally called the RRR motif^52^, present on 16 protomers of NUP153^17^.

Based on our data, the difference in binding affinities between the canonical FxFG motif and the enhancer-coupled PSGVFTFG super-motif of NUP153 to CLPs is approximately 1,000-fold (K_D_,_app_ ≥ 1.3 mM and about 1.5 μM respectively). This contributes to the asymmetry of various FG-NUP “signatures” that directly interact with HIV capsid cores along the cytoplasmic-nuclear axis of the NPC.

Furthermore, the relationship between the binding potential (strength) of FG/FxFG binding enhancers and their position within the zones of the NPC shows increasing binding potential toward the “nuclear basket.” We demonstrate that the affinity of these virus-adopted binding enhancers to capsid cores increases in the following order: NUP58 < POM121 < NUP153. The first two NUPs are members of the NPC core. While NUP58 lines the central transport conduit of the NPC^18,60,75–78^, POM121 is considered a core NUP due to its nature as a transmembrane protein. However, recent data show that POM121 is asymmetrically distributed at the NPC core toward the nucleoplasmic side, as revealed by super-resolution microscopy techniques^83,84^. This further supports the idea of a gradient of increasing affinities toward the nuclear periphery for the HIV capsid core.

Therefore, we suggest that the natural spatial arrangement of FG-NUPs with various FG repeats and enhancers regulates the affinity gradient of interactions between the capsid core and these signatures. FG motifs and enhancers are naturally distributed within the NPC (**Figure 1**) into distinct zones, forming an affinity gradient along the cytoplasmic-nuclear axis. We propose that this affinity gradient involves the sequential binding of the capsid core to different zones of FG-NUPs. The HIV core binds to FG-NUPs with progressively increasing affinity as it moves from cytoplasmic filaments (zone of FG/FxFG) to the core of the NPC (zone of GLFG of NUP98 and FG/FxFG enhancer(s) of NUP58 (POM121)) and the “nuclear basket” (zone of non-canonical FG super-motif of NUP153 and the enhancer(s) of NUP153 (POM121)) (**Figure 8**). This contrasts with the transport of cellular Kap-cargo complexes, which do not utilize the “FG” affinity gradient^85^. Recent findings show structural deformation and/or cracking of the NPC structural scaffold during the translocation of HIV capsid cores^9^. Moreover, the role of GLFG repeats was documented in stabilizing the NPC scaffold, acting as Velcro within the NPC architecture^86^. Our data indicate that GLFG motifs are the strongest binders to capsids among canonical FG-repeats. Therefore, we suggest that this interaction between GLFG motifs and capsid core destabilizes the Velcro, deforming/ripping the structural scaffold of the nuclear pore.

**Figure 8.**
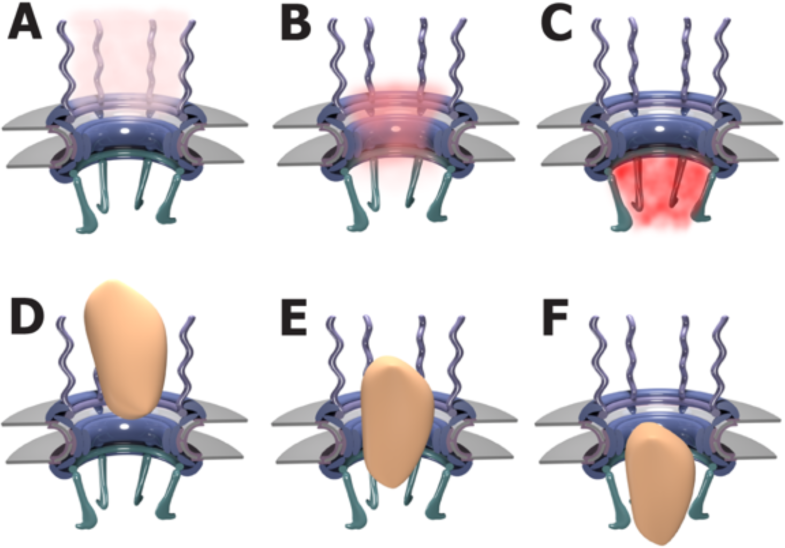
The model of HIV capsid core translocation through the NPC by affinity gradient. (A) Zone of FG/FxFG at the cytoplasmic side. (B) Zone of primarily NUP98 GLFG motifs and FG/FxFG binding enhancer(s) of NUP58 (POM121) at the core of the NPC. (C) FG/FxFG zone containing NUP153 FG super-motif and binding enhancer(s) of NUP153 (POM121) protruding to the “nuclear basket”. Increasing binding potentials of the zones are indicated by different colors from light (A) to medium (B) and dark red (C) respectively. The translocation of HIV capsid (orange) as passing from the cytoplasmic filaments (D) to the core of the NPC (E) and to the “nuclear basket” (F). For simplicity, only structured parts of the NPC cartoon are depicted.

Finally, after the HIV capsid core reaches the “nuclear basket,” the nuclear protein CPSF6 appears to facilitate the completion of nuclear import by releasing the capsid cores from the NPC^8,9,50^. CPSF6, similar to NUP153, contains another potent, non-canonical FG-containing motif PVLFPGQPFGQPPLG (CPSF6_313-327_)^47,87^, anchored around a central phenylalanine but otherwise engaging in diverse contacts with CA^45^. It has been suggested that CPSF6 competes with NUP153 FG super-motif for the same binding site on CA^44^, and by elevating the nuclear concentration of CPSF6, core-like particles speed up their intranuclear localization^88^.

In conclusion, it appears that HIV has evolved to take advantage of an asymmetric distribution of various flavors of nuclear pore complex FG motifs, binding enhancers, and non-canonical FG super-motifs. Our results comprise the first quantitative assessment of the individual interactions involved. As is often the case in biomolecular systems, the spatial organization of multiple weak-binding interactions achieves both a substantial difference in total binding energy, favoring the target destination while also avoiding kinetic trapping along the way. Finally, the complexity of NPC structure suggests that it would be difficult for the host to mutate in such a way to defeat this interaction-energy gradient, making HIV the apparent winner of this evolutionary arms race in a key function of its infection cycle.

## LIMITATION OF THE STUDY

The HIV capsid core possesses on average ∼1,500 FG-binding pockets. Although the accumulated strength of multivalent interactions could potentially substantially increase avidity, the multivalency of cellular Kap-FG NUP interactions has a much weaker effect on the overall affinity than expected as for other multivalent systems^66^. The extent of multivalency of FG NUP interactions on the overall affinity (i.e., avidity) to HIV capsid core, is currently unknown and awaits further investigation.

## ACKNOWLEDGEMENTS

We thank Elias Coutavas and Ilka Tona for critical reading of the manuscript and M.G. Finn (Georgia Institute of Technology) for valuable suggestions. We thank Joe Alexander for artwork. This research was supported by the National Institutes of Health AI176946 (I.M., S.G.S.), U54 AI170855 (S.G.S. and R.A.D.), R01 AI120860 and P30 AI050409 (S.G.S.). R.L.S. was supported in part by T32 AI157855. The content is solely the responsibility of the authors and does not necessarily represent the official views of the National Institutes of Health. S.G.S. acknowledges funding from the Nahmias-Schinazi Distinguished Chair in Research. This research used resources 17-ID-1 (AMX) or 17-ID-2 (FMX) of the National Synchrotron Light Source II; a U.S. Department of Energy (DOE) Office of Science User Facility operated for the DOE Office of Science by Brookhaven National Laboratory under Contract No. DE-SC0012704. The Center for BioMolecular Structure (CBMS) is primarily supported by the National Institutes of Health, National Institute of General Medical Sciences (NIGMS) through a Center Core P30 Grant (P30GM133893), and by the DOE Office of Biological and Environmental Research (KP1607011).

## AUTHOR CONTRIBUTIONS

Project conceptualization: I.M., S.G.S. Sample preparation and data acquisition: I.M., R.L.S., Z.C.L., A.E.C., K.A., J.S.W. Analysis: I.M., R.L.S., K.A.K., R.A.D., S.G.S. Writing, revising, editing: I.M., S.G.S. Methodology: I.M., R.L.S., K.A., K.A.K., R.A.D., S.G.S. Resources: I.M., S.G.S.

## METHODS

### Expression of proteins and peptides

Untagged HIV-1 CA_WT_ construct (in pET-28a) was expressed in *E. coli* BL21(DE3) at 37 °C in the presence of 1 mM isopropyl-thio-β-D-galactoside (IPTG), for 6 hrs. Untagged HIV-1 CA_A14C/E45C/W184A/M185A_ construct (in pET-11a) was expressed in *E. coli* BL21 Codon-Plus(DE3)-RIL at 25 °C in the presence of 1 mM IPTG, for 12 hrs.

Bivalent FG-, FSFG- and GLFG-derived constructs in pMAL-c5x (see Figure S2B) were expressed as N-terminally tagged MBP proteins at 30 °C in the presence of 0.2 mM IPTG, for 3 hrs. Bivalent GLFG-derived construct (in pET-28a) containing Cys residue (see Figure S2C) was expressed as N-terminally tagged 6 His-containing protein at 30 °C in the presence of 0.2 mM IPTG, for 3 hrs.

Monovalent, 6-Carboxyfluorescein (6-FAM) conjugated FG-, FSFG- and GLFG-containing peptides (see Figure S2A), unconjugated peptides used for crystallization (see Figure S2D), 6-FAM conjugated FTFG/FG-containing peptides of NUP153, POM121, NUP58 with and without their C-terminal basic residues, and 6-FAM conjugated C-terminal basic residues-containing peptides of NUP153, POM121, NUP58 (see Figure S8) were chemically synthesized and purchased from GenScript.

### Purification of CA proteins

Cell pellets of CA_A14C/E45C/W184A/M185A_ were resuspended in a lysis buffer (50 mM Tris-HCl, pH 8.0, 50 mM NaCl, 100 mM β-mercaptoethanol) supplemented with 12.5 mg/ml of lysozyme and incubated on ice for 1 hr. Cell lysate was prepared by sonication and clarified by centrifugation followed by filtration through 0.45 μm filter. CA proteins were then precipitated with saturated ammonium sulphate (in 2:1 ratio) for 1 hr on ice with occasional stirring. Precipitated proteins were pelleted by centrifugation and resuspended in a lysis buffer. After additional centrifugation, the supernatant was dialyzed against 25 mM Tris-HCl, pH 8.1, 20 mM β-mercaptoethanol, overnight at 4 °C. After dialysis, the sample was filtered through a 0.22 μm filter and loaded onto HiTrap Q XL column (Cytiva). CA proteins were eluted by a gradient of NaCl in dialysis buffer. Subsequently, the fractions of eluted CA were dialyzed against 25 mM MES, pH 5.5, 20 mM β-mercaptoethanol, clarified again by filtration, loaded onto HiTrap SP XL column (Cytiva) and eluted by a gradient of NaCl in dialysis buffer. Pooled fractions were dialyzed against storage buffer (20 mM Tris-HCl, pH 8.1, 40 mM NaCl) and concentrated.

Cell pellets of CA_WT_ were resuspended in a lysis buffer containing 20 mM Tris-HCl, pH 8.0, 1 mM Tris(2-carboxyethyl)phosphine (TCEP), 2 mM PMSF and lysed by sonication. The lysate was cleared by centrifugation and nucleic acids were precipitated from the supernatant by adding of polyethylenimine to a final concentration of 0.3%. After centrifugation, soluble proteins were precipitated using ammonium sulphate and collected by centrifugation. The pellet was resuspended in 50 mM Tris-HCl, pH 8.0, 1 mM TCEP and desalted using a HiPrep 26/10 column (Cytiva). The eluate was then subjected to tandem anion-cation exchange chromatography using HiTrap Q HP and HiTrap SP HP columns (Cytiva). The flow-through was collected and further purification was achieved with size exclusion chromatography (SEC) using a Superdex 75 Increase 10/300 GL column (Cytiva) equilibrated in a buffer containing 20 mM Tris-HCl, pH 8.0, 1 mM TCEP. The peak fractions were pooled, concentrated to 25 mg/ml, flash frozen in liquid N_2_ and stored in −80 °C until further analysis.

### Preparation of crosslinked CA hexamers

Concentrated CA_A14C/E45C/W184A/M185A_ proteins (≥15 mg/ml) were dialyzed in four subsequent steps against: 50 mM Tris-HCl, pH 8.1, 3 M NaCl, 50 mM β-mercaptoethanol (overnight), 50 mM Tris-HCl, pH 8.1, 3 M NaCl, 0.2 mM β-mercaptoethanol (8 hrs to overnight), 50 mM Tris-HCl, pH 8.1, 3 M NaCl (overnight), and 20 mM Tris-HCl, pH 8.1, 40 mM NaCl (overnight). All dialysis steps were performed at 4 °C. Dialyzed crosslinked CA hexamers were subjected to SEC. The proteins were run on HiLoad 26/600 Superdex 200 prep grade column (GE Healthcare), equilibrated in a buffer containing 20 mM Tris-HCl, pH 8.1, 40 mM NaCl. The peak fractions were pooled, concentrated to ∼160 mg/ml, flash frozen in liquid N_2_ and stored in −80 °C until further analysis.

### Purification of bivalent FG-peptides

Cell pellets containing MBP tagged FG-, FSFG- and GLFG-derived peptides were resuspended in 20 mM Tris-HCl, pH 8.0, 200 mM NaCl, 0.4 mM PMSF and lysed by sonication. After clarification by centrifugation, the supernatant was filtered through a 0.22 μm filter and loaded onto MBPTrap HP column (Cytiva). The proteins were eluted with 20 mM Tris-HCl, pH 8.0, 200 mM NaCl supplemented with 10 mM D-(+)-Maltose. MBP tag was cleaved by Factor Xa during dialysis against 50 mM Hepes, pH 8.0, 100 mM NaCl, overnight at 4 °C. The proteins were run on Superdex 200 Increase 10/300 GL column (Cytiva), equilibrated in the same buffer. The peak fractions containing cleaved peptides were pooled, flash frozen in liquid N_2_ and stored in −80 °C until further analysis.

Cell pellet with 6 His-fusion GLFG-derived peptide was resuspended in 25 mM Tris-HCl, pH 7.4, 250 mM NaCl, 5 mM imidazole, 1 mM TCEP and lysed by sonication. Cell lysate was clarified by centrifugation followed by filtration of the supernatant through 0.45 μm filter. The supernatant was then loaded onto HisTrap FF column (Cytiva) and the column was washed first with a buffer containing 25 mM Tris-HCl, pH 7.4, 250 mM NaCl, 5 mM imidazole, 1 mM TCEP and then with 25 mM Tris-HCl, pH 7.4, 250 mM NaCl and 1 mM TCEP. The bound peptide was immediately labelled by a column-based fluorochrome-conjugation approach (see below).

### Labeling of bivalent FG-peptides

The column-bound 6 His-fusion GLFG-derived peptide containing Cys residue (see Figure S2C) was labelled with Alexa Fluor 488 C5-maleimide (Invitrogen). In order to conjugate bound peptides with a fluorophore, the column was disconnected and ∼1.5 column volume of fluorophore solution was loaded by a syringe and the flow-through was passed again twice over the column. The solution contained 1 mM Alexa Fluor 488 C5-maleimide, 25 mM Tris-HCl, pH 7.4, 250 mM NaCl, 1 mM TCEP and a residual, ∼4.5% dimethyl sulfoxide from a fluorophore-dissolved stock solution. The column was reconnected and washed first with 25 mM Tris-HCl, pH 7.4, 250 mM NaCl, 1 mM TCEP, and then with the washing buffer without TCEP. Peptides were eluted by a gradient of imidazole supplemented in the same buffer. 6 His-tag was cleaved by thrombin during dialysis against 25 mM Tris-HCl, pH 7.4, 250 mM NaCl, overnight at 4 °C. The proteins were run on Superdex 200 Increase 10/300 GL column (Cytiva), equilibrated in the same buffer. The peak fractions containing cleaved and conjugated peptides were pooled, concentrated to ∼1 mM and flash frozen in liquid N_2_ and stored in −80 °C.

Concentrated, MBP cleaved and purified FG-, FSFG- and GLFG-derived peptides (see Figure S2B) were diluted to 5-10 mg/ml in 50 mM Hepes, pH 8.0, 100 mM NaCl and concentrated Alexa Fluor 647 NHS ester (Lumiprobe) was added in a molar ratio 1:10. The reaction was incubated overnight at 4 °C and unreacted fluorophores were quenched by addition of ethylenediamine, pH 8.0 to a final concentration of 50 mM. The conjugated peptides were then separated from unreacted fluorophores by SEC using Superdex 200 Increase 10/300 GL column (Cytiva) equilibrated in 50 mM Hepes, pH 8.0, 100 mM NaCl. The peak fractions of labeled peptides were pooled, concentrated to ∼1 mM and flash frozen in liquid N_2_ and stored in −80°C.

### Preparation of *E. coli* extract

BL21 Codon-Plus(DE3)-RIL cells were grown to an A_600_ of 0.8 and incubated for 16 hrs at 37 °C. Cell pellet was resuspended in 10 mM Tris-HCl, pH 8.0, 100 mM NaCl, 1 mM PMSF and lysed by a sonication. After the initial centrifugation, the supernatant was clarified by ultracentrifugation at 145,000 x g_av_ for 90 min at 4 °C. Protein concentration in the lysate was estimated by BCA assay to ∼9 mg/ml.

### Assembly of core-like particles (CLPs)

A buffer containing 50 mM MES, pH 6.2, 1 mM TCEP and 2 mM IP_6_ (Tokyo Chemical Company) was pre-warmed to 37 °C. Purified untagged CA_WT_ was then added to a final concentration of 400 μM and incubated at 37 °C for 30 min. Assemblies were stored at 4 °C and processed for MST assay withing 24 hours. Prior to the MST assay, CA lattice assemblies were screened by negative stain transmission electron microscopy (TEM) using Talos 120C TEM (see Figure S7).

### Microscale thermophoresis (MST)

MST experiments were performed and analyzed using a Monolith NT.115 (NanoTemper Technologies, Germany). To determine the affinity of monovalent FG-, FSFG- and GLFG-derived peptides to CA, 40 nM of 6-FAM labeled peptides were incubated with increasing concentrations of crosslinked CA hexamers, in 50 mM Tris-HCl, pH 8.1, 150 mM NaCl supplemented with 0.5 mg/ml bovine serum albumin (BSA) and loaded into NT.115 series standard capillaries (NanoTemper). 6-FAM fluorescence was monitored using nano-Blue channel (excitation 493 nm, emission 521 nm) with an excitation power of 20%, medium IR-laser power, and IR-laser on-time of 20 seconds. *n* = 2, 2, 3 technical replicates for FG, FSFG and GLFG respectively. The avidity of bivalent FG-, FSFG- and GLFG-derived peptides to CA was evaluated by incubation of 200 nM of Alexa Fluor 647 conjugated peptides with a range of concentrations of CA hexamers in 50 mM Tris-HCl, pH 8.1, 150 mM NaCl, 0.1% PEG 8,000. Samples were transferred to NT.115 series premium capillaries (NanoTemper). Alexa Fluor 647 fluorescence was monitored using nano-Red channel (excitation 650 nm, emission 670 nm) with an excitation power of 30%, 50% and 60% (for FG-, FSFG- and GLFG-peptides respectively), high IR-laser power, and IR-laser on-time of 5 seconds. *n* = 3 technical replicates for all three peptides. To assess the affinity of monovalent and bivalent GLFG-derived peptides to CA in the presence of weak binders, the reaction buffer (50 mM Tris-HCl, pH 8.1, 150 mM NaCl) was supplemented with 0.1 mg/ml *E.coli* lysate. 40 nM of 6-FAM labeled monovalent peptides or 500 nM of Alexa Fluor 488 conjugated bivalent peptides were incubated with increasing concentrations of CA hexamers and loaded into NT.115 series standard capillaries. 6-FAM fluorescence was monitored using nano-Blue channel with an excitation power of 50%, medium IR-laser power, and IR-laser on-time of 20 seconds. Alexa Fluor 488 fluorescence was monitored using nano-Blue channel with an excitation power of 20%, medium IR-laser power, and IR-laser on-time of 20 seconds. *n* = 2 technical replicates for bivalent peptides. The affinity of NUP153-, POM121- and NUP58-derived peptides to CLPs was assessed by incubation of 40 nM of 6-FAM conjugated peptides with a range of CLP concentrations in 50 mM MES, pH 6.2, 8.7 mM Tris-HCl, pH 8.0, 2 mM IP_6_, 1 mM TCEP and 1 mg/ml BSA. NT.115 series premium capillaries were used for the assay. For peptides of C-terminal basic residues, 6-FAM fluorescence was monitored in nano-Blue channel with an excitation power of 40%, 90% and 20% (NUP153, POM121 and NUP58 respectively), medium IR-laser power, and IR-laser on-time of 20 seconds. The experiments with POM121 peptide exhibited ligand-induced photobleaching and the scans were recorded in bleaching rate mode. *n* = 5, 3, 5 technical replicates for NUP153-Cterm, POM121-Cterm and NUP58-Cterm respectively. For FG/FTFG peptides with or without C-terminal basic residues, 6-FAM fluorescence was monitored in nano-Blue channel with an excitation power of 80%, medium IR-laser power, and IR-laser on-time of 20 seconds. The experiments with NUP153 and POM121 peptides exhibited ligand-induced photobleaching and the scans were recorded in bleaching rate mode. *n* = 4, 3, 3 technical replicates for NUP153-FTFG, POM121-FG and NUP58-FG respectively. *n* = 4, 3, 4 technical replicates for NUP153-FTFG•Cterm, POM121-FG•Cterm and NUP58-FG•Cterm respectively.

Technical replicates were performed from distinct samples. The data were analyzed with MO.AffinityAnalysis v2.3 (NanoTemper). The EC50 values were determined using a nonlinear regression fit of [inhibitor] vs. response with GraphPad Prism 10.2.3. For simplicity, we refer to EC50 values as K_D_ apparent (K_D_,_app_) in the text.

### Crystallization

∼0.07 mM of crosslinked CA hexamers (0.4 mM of crosslinked CA_A14C/E45C/W184A/M185A_ proteins) were initially co-crystallized with 4 mM of FG-, FSFG- and GLFG-derived peptides (see Figure S2D) using the hanging-drop method. The drops contained 1.3 μl of the protein/peptide (in 20 mM Tris-HCl, pH 8.1, 40 mM NaCl) and 1.3 μl of a reservoir solution consisting of 100 mM Hepes, pH 7.4 and 10% PEG 4,000. Orthorhombic crystals were grown at 18 °C within 7-10 days. For semi-quantitative analysis, we considered the final peptide concentration at the drop close to original concentration before mixing, i.e., 4 mM, due to a vapor diffusion. We therefore designate such concentration as an apparent concentration of the peptide in the text. To obtain higher apparent concentrations of corresponding peptides in the drop (∼9, ∼14, ∼34 and ∼43 mM respectively), the crystals were additionally soaked with higher concentrations of peptides dissolved in a reservoir solution. 0.6 μl of peptide (at 20, 36, 99.5 and 128 mM respectively) was added to ∼1.3 μl crystal drops and incubated for 2 days. For cryoprotection, the crystals were sequentially transferred into the reservoir solutions supplemented with the peptides of calculated apparent concentrations (see above) and the increasing amounts of glycerol up to a final concentration of 23% (v/v). Crystals were flash frozen in liquid N_2_.

### Data collection and structure determination

X-ray diffraction data were collected at the Brookhaven National Laboratory, NSLS-II, beamlines 17-ID-1 (AMX)^89^ and 17-ID-2 (FMX)^90^. X-ray intensities were processed using XDS^91^, and the CCP4 program suite (version 8.0.015)^92^ was used for subsequent calculations. The phase problem was solved using molecular replacement, with the coordinates of hexameric CA (PDB ID: 3H4E) as a search model. Initial phases were solved *via* PHASER (version 2.8.3)^93^. Several rounds of iterative model building and refinement were carried out using Coot (version 0.9.8.91)^94^, PHENIX (version 1.21-5207-000)^95^ and REFMAC (version 5.8.0419)^96^ respectively. Structure validation of final models was performed with MOLPROBITY (version 4.5.2)^97^. RMSD values were calculated using VMD (version 1.9.4)^98^. The figures showing structural information were generated in PyMOL (version 2.1.1)^99^ and UCSF ChimeraX (version 1.6.1)^100^. Coordinates and structure factors have been deposited in the RCSB Protein Data Bank (PDB). Data collection and refinement statistics are provided in Table S1.

## REFERENCES

1. Briggs, J. A., Wilk, T., Welker, R., Kräusslich, H. G., and Fuller, S. D. (2003). Structural organization of authentic, mature HIV-1 virions and cores. The EMBO journal, 22, 1707–1715. 10.1093/emboj/cdg143

2. Christensen, D. E., Ganser-Pornillos, B. K., Johnson, J. S., Pornillos, O., and Sundquist, W. I. (2020). Reconstitution and visualization of HIV-1 capsid-dependent replication and integration in vitro. Science, 370, eabc8420. 10.1126/science.abc8420

3. Engelman A. N. (2021). HIV capsid and integration targeting. Viruses, 13, 125. 10.3390/v13010125

4. Campbell, E. M., and Hope, T. J. (2015). HIV-1 capsid: the multifaceted key player in HIV-1 infection. Nature reviews. Microbiology, 13, 471–483. 10.1038/nrmicro3503

5. Matreyek, K. A., and Engelman, A. (2013). Viral and cellular requirements for the nuclear entry of retroviral preintegration nucleoprotein complexes. Viruses, 5, 2483–2511. 10.3390/v5102483

6. Lusic, M., and Siliciano, R. F. (2017). Nuclear landscape of HIV-1 infection and integration. Nature reviews. Microbiology, 15, 69–82. 10.1038/nrmicro.2016.162

7. Francis, A. C., and Melikyan, G. B. (2018). Live-cell imaging of early steps of single HIV-1 infection. Viruses, 10, 275. 10.3390/v10050275

8. Zila, V., Margiotta, E., Turoňová, B., Müller, T. G., Zimmerli, C. E., Mattei, S., Allegretti, M., Börner, K., Rada, J., Müller, B., Lusic, M., Kräusslich, H. G., and Beck, M. (2021). Cone-shaped HIV-1 capsids are transported through intact nuclear pores. Cell, 184, 1032–1046.e18. 10.1016/j.cell.2021.01.025

9. Kreysing, J. P., Heidari, M., Zila, V., Cruz-León, S., Obarska-Kosinska, A., Laketa, V., Rohleder, L., Welsch, S., Köfinger, J., Turoňová, B., Hummer, G., Kräusslich, H. G., and Beck, M. (2025). Passage of the HIV capsid cracks the nuclear pore. Cell, 188, 930–943.e21. 10.1016/j.cell.2024.12.008

10. Hou, Z., Shen, Y., Fronik, S., Shen, J., Shi, J., Xu, J., Chen, L., Hardenbrook, N., Engelman, A. N., Aiken, C., and Zhang, P. (2025). HIV-1 nuclear import is selective and depends on both capsid elasticity and nuclear pore adaptability. Nature microbiology, 10, 1868–1885. 10.1038/s41564-025-02054-z

11. Hou, Z., Fronik, S., Shen, Y., Chen, L., Thompson, C., Neumann, S., and Zhang, P. (2025). Direct visualization of HIV-1 core nuclear import and its interplay with the nuclear pore. EMBO reports, 10.1038/s44319-025-00567-6. Advance online publication. 10.1038/s44319-025-00567-6

12. Burdick, R. C., Li, C., Munshi, M., Rawson, J., Nagashima, K., Hu, W. S., and Pathak, V. K. (2020). HIV-1 uncoats in the nucleus near sites of integration. Proceedings of the National Academy of Sciences of the United States of America, 117, 5486–5493. 10.1073/pnas.1920631117

13. Li, C., Burdick, R. C., Nagashima, K., Hu, W. S., and Pathak, V. K. (2021). HIV-1 cores retain their integrity until minutes before uncoating in the nucleus. Proceedings of the National Academy of Sciences of the United States of America, 118, e2019467118. 10.1073/pnas.2019467118

14. Cronshaw, J. M., Krutchinsky, A. N., Zhang, W., Chait, B. T., and Matunis, M. J. (2002). Proteomic analysis of the mammalian nuclear pore complex. The Journal of cell biology, 158, 915–927. 10.1083/jcb.200206106

15. Gall J. G. (1967). Octagonal nuclear pores. The Journal of cell biology, 32, 391–399. 10.1083/jcb.32.2.391

16. Unwin, P. N., and Milligan, R. A. (1982). A large particle associated with the perimeter of the nuclear pore complex. The Journal of cell biology, 93, 63–75. 10.1083/jcb.93.1.63

17. Singh, D., Soni, N., Hutchings, J., Echeverria, I., Shaikh, F., Duquette, M., Suslov, S., Li, Z., van Eeuwen, T., Molloy, K., Shi, Y., Wang, J., Guo, Q., Chait, B. T., Fernandez-Martinez, J., Rout, M. P., Sali, A., and Villa, E. (2024). The molecular architecture of the nuclear basket. Cell, 187, 5267–5281.e13. 10.1016/j.cell.2024.07.020

18. Schuller, A. P., Wojtynek, M., Mankus, D., Tatli, M., Kronenberg-Tenga, R., Regmi, S. G., Dip, P. V., Lytton-Jean, A., Brignole, E. J., Dasso, M., Weis, K., Medalia, O., and Schwartz, T. U. (2021). The cellular environment shapes the nuclear pore complex architecture. Nature, 598, 667–671. 10.1038/s41586-021-03985-3

19. Zimmerli, C. E., Allegretti, M., Rantos, V., Goetz, S. K., Obarska-Kosinska, A., Zagoriy, I., Halavatyi, A., Hummer, G., Mahamid, J., Kosinski, J., and Beck, M. (2021). Nuclear pores dilate and constrict in cellulo. Science, 374, eabd9776. 10.1126/science.abd9776

20. Rout, M. P., Aitchison, J. D., Suprapto, A., Hjertaas, K., Zhao, Y., and Chait, B. T. (2000). The yeast nuclear pore complex: composition, architecture, and transport mechanism. The Journal of cell biology, 148, 635–651. 10.1083/jcb.148.4.635

21. Yang W. (2011). ’Natively unfolded’ nucleoporins in nucleocytoplasmic transport: clustered or evenly distributed? Nucleus, 2, 10–16. 10.4161/nucl.2.1.13818

22. Wente, S. R., Rout, M. P., and Blobel, G. (1992). A new family of yeast nuclear pore complex proteins. The Journal of cell biology, 119, 705–723. 10.1083/jcb.119.4.705

23. Radu, A., Moore, M. S., and Blobel, G. (1995). The peptide repeat domain of nucleoporin Nup98 functions as a docking site in transport across the nuclear pore complex. Cell, 81, 215–222. 10.1016/0092-8674(95)90331-3

24. Mohr, D., Frey, S., Fischer, T., Güttler, T., and Görlich, D. (2009). Characterisation of the passive permeability barrier of nuclear pore complexes. The EMBO journal, 28, 2541–2553. 10.1038/emboj.2009.200

25. Allen, N. P., Huang, L., Burlingame, A., and Rexach, M. (2001). Proteomic analysis of nucleoporin interacting proteins. The Journal of biological chemistry, 276, 29268–29274. 10.1074/jbc.M102629200

26. Kalita, J., Kapinos, L. E., and Lim, R. (2021). On the asymmetric partitioning of nucleocytoplasmic transport - recent insights and open questions. Journal of cell science, 134, jcs240382. 10.1242/jcs.240382

27. Floer, M., Blobel, G., and Rexach, M. (1997). Disassembly of RanGTP-karyopherin beta complex, an intermediate in nuclear protein import. The Journal of biological chemistry, 272, 19538–19546. 10.1074/jbc.272.31.19538

28. Macara I. G. (2001). Transport into and out of the nucleus. Microbiology and molecular biology reviews : MMBR, 65, 570–594. 10.1128/MMBR.65.4.570-594.2001

29. Izaurralde, E., Kutay, U., von Kobbe, C., Mattaj, I. W., and Görlich, D. (1997). The asymmetric distribution of the constituents of the Ran system is essential for transport into and out of the nucleus. The EMBO journal, 16, 6535–6547. 10.1093/emboj/16.21.6535

30. Rexach, M., and Blobel, G. (1995). Protein import into nuclei: association and dissociation reactions involving transport substrate, transport factors, and nucleoporins. Cell, 83, 683–692. 10.1016/0092-8674(95)90181-7

31. Kalita, J., Kapinos, L. E., and Lim, R. (2021). On the asymmetric partitioning of nucleocytoplasmic transport - recent insights and open questions. Journal of cell science, 134, jcs240382. 10.1242/jcs.240382

32. Chook, Y. M., and Süel, K. E. (2011). Nuclear import by karyopherin-βs: recognition and inhibition. Biochimica et biophysica acta, 1813, 1593–1606. 10.1016/j.bbamcr.2010.10.014

33. Matsuura Y. (2016). Mechanistic insights from structural analyses of Ran-GTPase-driven nuclear export of proteins and RNAs. Journal of molecular biology, 428, 2025–2039. 10.1016/j.jmb.2015.09.025

34. Wing, C. E., Fung, H. Y. J., and Chook, Y. M. (2022). Karyopherin-mediated nucleocytoplasmic transport. Nature reviews. Molecular cell biology, 23, 307–328. 10.1038/s41580-021-00446-7

35. Bayliss, R., Littlewood, T., Strawn, L. A., Wente, S. R., and Stewart, M. (2002). GLFG and FxFG nucleoporins bind to overlapping sites on importin-beta. The Journal of biological chemistry, 277, 50597–50606. 10.1074/jbc.M209037200

36. Bayliss, R., Littlewood, T., and Stewart, M. (2000). Structural basis for the interaction between FxFG nucleoporin repeats and importin-beta in nuclear trafficking. Cell, 102, 99–108. 10.1016/s0092-8674(00)00014-3

37. Fribourg, S., Braun, I. C., Izaurralde, E., and Conti, E. (2001). Structural basis for the recognition of a nucleoporin FG repeat by the NTF2-like domain of the TAP/p15 mRNA nuclear export factor. Molecular cell, 8, 645–656. 10.1016/s1097-2765(01)00348-3

38. Hough, L. E., Dutta, K., Sparks, S., Temel, D. B., Kamal, A., Tetenbaum-Novatt, J., Rout, M. P., and Cowburn, D. (2015). The molecular mechanism of nuclear transport revealed by atomic-scale measurements. eLife, 4, e10027. 10.7554/eLife.10027

39. Milles, S., Mercadante, D., Aramburu, I. V., Jensen, M. R., Banterle, N., Koehler, C., Tyagi, S., Clarke, J., Shammas, S. L., Blackledge, M., Gräter, F., and Lemke, E. A. (2015). Plasticity of an ultrafast interaction between nucleoporins and nuclear transport receptors. Cell, 163, 734–745. 10.1016/j.cell.2015.09.047

40. Raveh, B., Karp, J. M., Sparks, S., Dutta, K., Rout, M. P., Sali, A., and Cowburn, D. (2016). Slide-and-exchange mechanism for rapid and selective transport through the nuclear pore complex. Proceedings of the National Academy of Sciences of the United States of America, 113, 2489–2497. 10.1073/pnas.1522663113

41. Mattei, S., Glass, B., Hagen, W. J., Kräusslich, H. G., and Briggs, J. A. (2016). The structure and flexibility of conical HIV-1 capsids determined within intact virions. Science, 354, 1434–1437. 10.1126/science.aah4972

42. Dickson, C. F., Hertel, S., Tuckwell, A. J., Li, N., Ruan, J., Al-Izzi, S. C., Ariotti, N., Sierecki, E., Gambin, Y., Morris, R. G., Towers, G. J., Böcking, T., and Jacques, D. A. (2024). The HIV capsid mimics karyopherin engagement of FG-nucleoporins. Nature, 626, 836–842. 10.1038/s41586-023-06969-7

43. Fu, L., Weiskopf, E. N., Akkermans, O., Swanson, N. A., Cheng, S., Schwartz, T. U., and Görlich, D. (2024). HIV-1 capsids enter the FG phase of nuclear pores like a transport receptor. Nature, 626, 843–851. 10.1038/s41586-023-06966-w

44. Matreyek, K. A., Yücel, S. S., Li, X., and Engelman, A. (2013). Nucleoporin NUP153 phenylalanine-glycine motifs engage a common binding pocket within the HIV-1 capsid protein to mediate lentiviral infectivity. PLoS pathogens, 9, e1003693. 10.1371/journal.ppat.1003693

45. Price, A. J., Jacques, D. A., McEwan, W. A., Fletcher, A. J., Essig, S., Chin, J. W., Halambage, U. D., Aiken, C., and James, L. C. (2014). Host cofactors and pharmacologic ligands share an essential interface in HIV-1 capsid that is lost upon disassembly. PLoS pathogens, 10, e1004459. 10.1371/journal.ppat.1004459

46. Gres, A. T., Kirby, K. A., McFadden, W. M., Du, H., Liu, D., Xu, C., Bryer, A. J., Perilla, J. R., Shi, J., Aiken, C., Fu, X., Zhang, P., Francis, A. C., Melikyan, G. B., and Sarafianos, S. G. (2023). Multidisciplinary studies with mutated HIV-1 capsid proteins reveal structural mechanisms of lattice stabilization. Nature communications, 14, 5614. 10.1038/s41467-023-41197-7

47. Price, A. J., Fletcher, A. J., Schaller, T., Elliott, T., Lee, K., KewalRamani, V. N., Chin, J. W., Towers, G. J., and James, L. C. (2012). CPSF6 defines a conserved capsid interface that modulates HIV-1 replication. PLoS pathogens, 8, e1002896. 10.1371/journal.ppat.1002896

48. Bhattacharya, A., Alam, S. L., Fricke, T., Zadrozny, K., Sedzicki, J., Taylor, A. B., Demeler, B., Pornillos, O., Ganser-Pornillos, B. K., Diaz-Griffero, F., Ivanov, D. N., and Yeager, M. (2014). Structural basis of HIV-1 capsid recognition by PF74 and CPSF6. Proceedings of the National Academy of Sciences of the United States of America, 111, 18625–18630. 10.1073/pnas.1419945112

49. Achuthan, V., Perreira, J. M., Sowd, G. A., Puray-Chavez, M., McDougall, W. M., Paulucci-Holthauzen, A., Wu, X., Fadel, H. J., Poeschla, E. M., Multani, A. S., Hughes, S. H., Sarafianos, S. G., Brass, A. L., and Engelman, A. N. (2018). Capsid-CPSF6 interaction licenses nuclear HIV-1 trafficking to sites of viral DNA integration. Cell host & microbe, 24, 392–404.e8. 10.1016/j.chom.2018.08.002

50. Bejarano, D. A., Peng, K., Laketa, V., Börner, K., Jost, K. L., Lucic, B., Glass, B., Lusic, M., Müller, B., and Kräusslich, H. G. (2019). HIV-1 nuclear import in macrophages is regulated by CPSF6-capsid interactions at the nuclear pore complex. eLife, 8, e41800. 10.7554/eLife.41800

51. Chaudhuri, E., Jang, S., Chakraborty, R., Radhakrishnan, R., Arnarson, B., Prakash, P., Cornish, D., Rohlfes, N., Singh, P. K., Shi, J., Aiken, C., Campbell, E., Hultquist, J., Balsubramaniam, M., Engelman, A. N., and Dash, C. (2025). CPSF6 promotes HIV-1 preintegration complex function. Journal of virology, e0049025. 10.1128/jvi.00490-25

52. Shen, Q., Kumari, S., Xu, C., Jang, S., Shi, J., Burdick, R. C., Levintov, L., Xiong, Q., Wu, C., Devarkar, S. C., Tian, T., Tripler, T. N., Hu, Y., Yuan, S., Temple, J., Feng, Q., Lusk, C. P., Aiken, C., Engelman, A. N., Perilla, J. R., … Xiong, Y. (2023). The capsid lattice engages a bipartite NUP153 motif to mediate nuclear entry of HIV-1 cores. Proceedings of the National Academy of Sciences of the United States of America, 120, e2202815120. 10.1073/pnas.2202815120

53. Shen, Q., Wu, C., Freniere, C., Tripler, T. N., and Xiong, Y. (2021). Nuclear Import of HIV-1. Viruses, 13, 2242. 10.3390/v13112242

54. Aramburu, I. V., and Lemke, E. A. (2017). Floppy but not sloppy: Interaction mechanism of FG-nucleoporins and nuclear transport receptors. Seminars in cell & developmental biology, 68, 34–41. 10.1016/j.semcdb.2017.06.026

55. Reichelt, R., Holzenburg, A., Buhle, E. L., Jr, Jarnik, M., Engel, A., and Aebi, U. (1990). Correlation between structure and mass distribution of the nuclear pore complex and of distinct pore complex components. The Journal of cell biology, 110, 883–894. 10.1083/jcb.110.4.883

56. Huang, K., Tagliazucchi, M., Park, S. H., Rabin, Y., and Szleifer, I. (2020). Nanocompartmentalization of the nuclear pore lumen. Biophysical journal, 118, 219–231. 10.1016/j.bpj.2019.11.024

57. Kim, S. J., Fernandez-Martinez, J., Nudelman, I., Shi, Y., Zhang, W., Raveh, B., Herricks, T., Slaughter, B. D., Hogan, J. A., Upla, P., Chemmama, I. E., Pellarin, R., Echeverria, I., Shivaraju, M., Chaudhury, A. S., Wang, J., Williams, R., Unruh, J. R., Greenberg, C. H., Jacobs, E. Y., … Rout, M. P. (2018). Integrative structure and functional anatomy of a nuclear pore complex. Nature, 555, 475–482. 10.1038/nature26003

58. Patel, S. S., Belmont, B. J., Sante, J. M., and Rexach, M. F. (2007). Natively unfolded nucleoporins gate protein diffusion across the nuclear pore complex. Cell, 129, 83–96. 10.1016/j.cell.2007.01.044

59. Ori, A., Banterle, N., Iskar, M., Andrés-Pons, A., Escher, C., Khanh Bui, H., Sparks, L., Solis-Mezarino, V., Rinner, O., Bork, P., Lemke, E. A., and Beck, M. (2013). Cell type-specific nuclear pores: a case in point for context-dependent stoichiometry of molecular machines. Molecular systems biology, 9, 648. 10.1038/msb.2013.4

60. Petrovic, S., Samanta, D., Perriches, T., Bley, C. J., Thierbach, K., Brown, B., Nie, S., Mobbs, G. W., Stevens, T. A., Liu, X., Tomaleri, G. P., Schaus, L., and Hoelz, A. (2022). Architecture of the linker-scaffold in the nuclear pore. Science, 376, eabm9798. 10.1126/science.abm9798

61. Ma, J., Kelich, J. M., Junod, S. L., and Yang, W. (2017). Super-resolution mapping of scaffold nucleoporins in the nuclear pore complex. Journal of cell science, 130, 1299–1306. 10.1242/jcs.193912

62. Otsuka, S., Tempkin, J. O. B., Zhang, W., Politi, A. Z., Rybina, A., Hossain, M. J., Kueblbeck, M., Callegari, A., Koch, B., Morero, N. R., Sali, A., and Ellenberg, J. (2023). A quantitative map of nuclear pore assembly reveals two distinct mechanisms. Nature, 613, 575–581. 10.1038/s41586-022-05528-w

63. Denning, D. P., and Rexach, M. F. (2007). Rapid evolution exposes the boundaries of domain structure and function in natively unfolded FG nucleoporins. Molecular & cellular proteomics : MCP, 6, 272–282. 10.1074/mcp.M600309-MCP200

64. Hülsmann, B. B., Labokha, A. A., and Görlich, D. (2012). The permeability of reconstituted nuclear pores provides direct evidence for the selective phase model. Cell, 150, 738–751. 10.1016/j.cell.2012.07.019

65. Pornillos, O., Ganser-Pornillos, B. K., Banumathi, S., Hua, Y., and Yeager, M. (2010). Disulfide bond stabilization of the hexameric capsomer of human immunodeficiency virus. Journal of molecular biology, 401, 985–995. 10.1016/j.jmb.2010.06.042

66. Hayama, R., Sparks, S., Hecht, L. M., Dutta, K., Karp, J. M., Cabana, C. M., Rout, M. P., and Cowburn, D. (2018). Thermodynamic characterization of the multivalent interactions underlying rapid and selective translocation through the nuclear pore complex. The Journal of biological chemistry, 293, 4555–4563. 10.1074/jbc.AC117.001649

67. Tetenbaum-Novatt, J., Hough, L. E., Mironska, R., McKenney, A. S., and Rout, M. P. (2012). Nucleocytoplasmic transport: a role for nonspecific competition in karyopherin-nucleoporin interactions. Molecular & cellular proteomics : MCP, 11, 31–46. 10.1074/mcp.M111.013656

68. Bayliss, R., Littlewood, T., Strawn, L. A., Wente, S. R., and Stewart, M. (2002). GLFG and FxFG nucleoporins bind to overlapping sites on importin-beta. The Journal of biological chemistry, 277, 50597–50606. 10.1074/jbc.M209037200

69. Bayliss, R., Littlewood, T., and Stewart, M. (2000). Structural basis for the interaction between FxFG nucleoporin repeats and importin-beta in nuclear trafficking. Cell, 102, 99–108. 10.1016/s0092-8674(00)00014-3

70. Highland, C. M., Tan, A., Ricaña, C. L., Briggs, J. A. G., and Dick, R. A. (2023). Structural insights into HIV-1 polyanion-dependent capsid lattice formation revealed by single particle cryo-EM. Proceedings of the National Academy of Sciences of the United States of America, 120, e2220545120. 10.1073/pnas.2220545120

71. Erlendsson, S., and Teilum, K. (2021). Binding revisited-Avidity in cellular function and signaling. Frontiers in molecular biosciences, 7, 615565. 10.3389/fmolb.2020.615565

72. Cardarelli, F., Lanzano, L., and Gratton, E. (2011). Fluorescence correlation spectroscopy of intact nuclear pore complexes. Biophysical journal, 101, 27–29. 10.1016/j.bpj.2011.04.057

73. Stewart M. (2007). Molecular mechanism of the nuclear protein import cycle. Nature reviews. Molecular cell biology, 8, 195–208. 10.1038/nrm2114

74. Bley, C. J., Nie, S., Mobbs, G. W., Petrovic, S., Gres, A. T., Liu, X., Mukherjee, S., Harvey, S., Huber, F. M., Lin, D. H., Brown, B., Tang, A. W., Rundlet, E. J., Correia, A. R., Chen, S., Regmi, S. G., Stevens, T. A., Jette, C. A., Dasso, M., Patke, A., … Hoelz, A. (2022). Architecture of the cytoplasmic face of the nuclear pore. Science, 376, eabm9129. 10.1126/science.abm9129

75. Lin, D. H., Stuwe, T., Schilbach, S., Rundlet, E. J., Perriches, T., Mobbs, G., Fan, Y., Thierbach, K., Huber, F. M., Collins, L. N., Davenport, A. M., Jeon, Y. E., and Hoelz, A. (2016). Architecture of the symmetric core of the nuclear pore. Science, 352, aaf1015. 10.1126/science.aaf1015

76. von Appen, A., Kosinski, J., Sparks, L., Ori, A., DiGuilio, A. L., Vollmer, B., Mackmull, M. T., Banterle, N., Parca, L., Kastritis, P., Buczak, K., Mosalaganti, S., Hagen, W., Andres-Pons, A., Lemke, E. A., Bork, P., Antonin, W., Glavy, J. S., Bui, K. H., and Beck, M. (2015). In situ structural analysis of the human nuclear pore complex. Nature, 526, 140–143. 10.1038/nature15381

77. Kosinski, J., Mosalaganti, S., von Appen, A., Teimer, R., DiGuilio, A. L., Wan, W., Bui, K. H., Hagen, W. J., Briggs, J. A., Glavy, J. S., Hurt, E., and Beck, M. (2016). Molecular architecture of the inner ring scaffold of the human nuclear pore complex. Science, 352, 363–365. 10.1126/science.aaf0643

78. Mosalaganti, S., Obarska-Kosinska, A., Siggel, M., Taniguchi, R., Turoňová, B., Zimmerli, C. E., Buczak, K., Schmidt, F. H., Margiotta, E., Mackmull, M. T., Hagen, W. J. H., Hummer, G., Kosinski, J., and Beck, M. (2022). AI-based structure prediction empowers integrative structural analysis of human nuclear pores. Science, 376, eabm9506. 10.1126/science.abm9506

79. Bley, C. J., Nie, S., Mobbs, G. W., Petrovic, S., Gres, A. T., Liu, X., Mukherjee, S., Harvey, S., Huber, F. M., Lin, D. H., Brown, B., Tang, A. W., Rundlet, E. J., Correia, A. R., Chen, S., Regmi, S. G., Stevens, T. A., Jette, C. A., Dasso, M., Patke, A., … Hoelz, A. (2022). Architecture of the cytoplasmic face of the nuclear pore. Science, 376, eabm9129. 10.1126/science.abm9129

80. Fontana, P., Dong, Y., Pi, X., Tong, A. B., Hecksel, C. W., Wang, L., Fu, T. M., Bustamante, C., and Wu, H. (2022). Structure of cytoplasmic ring of nuclear pore complex by integrative cryo-EM and AlphaFold. Science, 376, eabm9326. 10.1126/science.abm9326

81. Sukegawa, J., and Blobel, G. (1993). A nuclear pore complex protein that contains zinc finger motifs, binds DNA, and faces the nucleoplasm. Cell, 72, 29–38. 10.1016/0092-8674(93)90047-t

82. Guan, T., Kehlenbach, R. H., Schirmer, E. C., Kehlenbach, A., Fan, F., Clurman, B. E., Arnheim, N., and Gerace, L. (2000). Nup50, a nucleoplasmically oriented nucleoporin with a role in nuclear protein export. Molecular and cellular biology, 20, 5619–5630. 10.1128/MCB.20.15.5619-5630.2000

83. Morgan, K. J., Carley, E., Coyne, A. N., Rothstein, J. D., Lusk, C. P., and King, M. C. (2025). Visualizing nuclear pore complex plasticity with pan-expansion microscopy. The Journal of cell biology, 224, e202409120. 10.1083/jcb.202409120

84. Andronov, L., Genthial, R., Hentsch, D., and Klaholz, B. P. (2022). splitSMLM, a spectral demixing method for high-precision multi-color localization microscopy applied to nuclear pore complexes. Communications biology, 5, 1100. 10.1038/s42003-022-04040-1

85. Zeitler, B., and Weis, K. (2004). The FG-repeat asymmetry of the nuclear pore complex is dispensable for bulk nucleocytoplasmic transport in vivo. The Journal of cell biology, 167, 583–590. 10.1083/jcb.200407156

86. Onischenko, E., Tang, J. H., Andersen, K. R., Knockenhauer, K. E., Vallotton, P., Derrer, C. P., Kralt, A., Mugler, C. F., Chan, L. Y., Schwartz, T. U., and Weis, K. (2017). Natively unfolded FG repeats stabilize the structure of the nuclear pore complex. Cell, 171, 904–917.e19. 10.1016/j.cell.2017.09.033

87. Lee, K., Mulky, A., Yuen, W., Martin, T. D., Meyerson, N. R., Choi, L., Yu, H., Sawyer, S. L., and Kewalramani, V. N. (2012). HIV-1 capsid-targeting domain of cleavage and polyadenylation specificity factor 6. Journal of virology, 86, 3851–3860. 10.1128/JVI.06607-11

88. Fu, L., Riedel, D., Kopecny, L., Schuh, M., and Görlich, D. (2025). Governed by surface amino acid composition: HIV capsid passage through the NPC barrier. bioRxiv : the preprint server for biology, 2025.03.13.643050. 10.1101/2025.03.13.643050

89. Schneider, D. K., Soares, A. S., Lazo, E. O., Kreitler, D. F., Qian, K., Fuchs, M. R., Bhogadi, D. K., Antonelli, S., Myers, S. S., Martins, B. S., Skinner, J. M., Aishima, J., Bernstein, H. J., Langdon, T., Lara, J., Petkus, R., Cowan, M., Flaks, L., Smith, T., Shea-McCarthy, G., … Jakoncic, J. (2022). AMX - the highly automated macromolecular crystallography (17-ID-1) beamline at the NSLS-II. Journal of synchrotron radiation, 29, 1480–1494. 10.1107/S1600577522009377

90. Schneider, D. K., Shi, W., Andi, B., Jakoncic, J., Gao, Y., Bhogadi, D. K., Myers, S. F., Martins, B., Skinner, J. M., Aishima, J., Qian, K., Bernstein, H. J., Lazo, E. O., Langdon, T., Lara, J., Shea-McCarthy, G., Idir, M., Huang, L., Chubar, O., Sweet, R. M., … Fuchs, M. R. (2021). FMX - the frontier microfocusing macromolecular crystallography beamline at the National Synchrotron Light Source II. Journal of synchrotron radiation, 28, 650–665. 10.1107/S1600577520016173

91. Kabsch W. (2010). XDS. Acta crystallographica. Section D, Biological crystallography, 66, 125–132. 10.1107/S0907444909047337

92. Agirre, J., Atanasova, M., Bagdonas, H., Ballard, C. B., Baslé, A., Beilsten-Edmands, J., Borges, R. J., Brown, D. G., Burgos-Mármol, J. J., Berrisford, J. M., Bond, P. S., Caballero, I., Catapano, L., Chojnowski, G., Cook, A. G., Cowtan, K. D., Croll, T. I., Debreczeni, J. É., Devenish, N. E., Dodson, E. J., … Yamashita, K. (2023). The CCP4 suite: integrative software for macromolecular crystallography. Acta crystallographica. Section D, Structural biology, 79, 449–461. 10.1107/S2059798323003595

93. McCoy, A. J., Grosse-Kunstleve, R. W., Adams, P. D., Winn, M. D., Storoni, L. C., and Read, R. J. (2007). Phaser crystallographic software. Journal of applied crystallography, 40, 658–674. 10.1107/S0021889807021206

94. Emsley, P., Lohkamp, B., Scott, W. G., and Cowtan, K. (2010). Features and development of Coot. Acta crystallographica. Section D, Biological crystallography, 66, 486–501. 10.1107/S0907444910007493

95. Afonine, P. V., Grosse-Kunstleve, R. W., Echols, N., Headd, J. J., Moriarty, N. W., Mustyakimov, M., Terwilliger, T. C., Urzhumtsev, A., Zwart, P. H., and Adams, P. D. (2012). Towards automated crystallographic structure refinement with phenix.refine. Acta crystallographica. Section D, Biological crystallography, 68, 352–367. 10.1107/S0907444912001308

96. Kovalevskiy, O., Nicholls, R. A., Long, F., Carlon, A., and Murshudov, G. N. (2018). Overview of refinement procedures within REFMAC5: utilizing data from different sources. Acta crystallographica. Section D, Structural biology, 74, 215–227. 10.1107/S2059798318000979

97. Williams, C. J., Headd, J. J., Moriarty, N. W., Prisant, M. G., Videau, L. L., Deis, L. N., Verma, V., Keedy, D. A., Hintze, B. J., Chen, V. B., Jain, S., Lewis, S. M., Arendall, W. B., 3rd, Snoeyink, J., Adams, P. D., Lovell, S. C., Richardson, J. S., and Richardson, D. C. (2018). MolProbity: More and better reference data for improved all-atom structure validation. Protein science, 27, 293–315. 10.1002/pro.3330

98. Humphrey, W., Dalke, A., and Schulten, K. (1996). VMD: visual molecular dynamics. Journal of molecular graphics, 14, 33–28. 10.1016/0263-7855(96)00018-5

99. DeLano, W. L. www.pymol.org (2002).

100. Pettersen, E. F., Goddard, T. D., Huang, C. C., Meng, E. C., Couch, G. S., Croll, T. I., Morris, J. H., and Ferrin, T. E. (2021). UCSF ChimeraX: Structure visualization for researchers, educators, and developers. Protein science, 30, 70–82. 10.1002/pro.3943

