## Supplement for "Translocation of HIV capsid core through the Nuclear Pore Complex by affinity gradient"

SUPPLEMENTAL INFORMATION

**Figure S1. The design of "universal" linker for various FG motifs**
(A) The sequence of two native, 14-mer "parental" human NUP98 FG-containing motifs (in red), GLFG (top) and FSFG (bottom). The residues above the primary sequences represent variable amino acid substitutions found in vertebrate family of NUP98 by sequence conservation analysis. (B) The inferred 14-mer sequences of "universal" peptides with embedded GLFG, FSFG, and FG motifs (red).

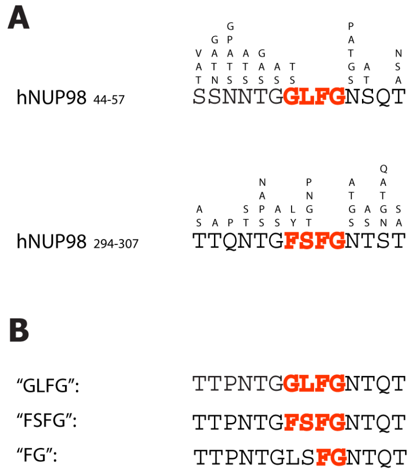

**Figure S2. The design of peptides with monovalent and bivalent FG motifs**
(A) Monovalent 6-Carboxyfluorescein (6-FAM) conjugated FG-, FSFG- and GLFG-containing peptides. (B) Two 14-mer sequences of "universal" GLFG-, FSFG- or FG- derived peptides (red) separated by unstructured, Gly-Ser rich sequences. The residual vector sequence is shown in italics. (C) Same as (B) except Ser-to-Cys mutation at the N-terminus. Residues conjugated with a fluorochrome are indicated (asterisk). (D) Unconjugated peptides used for crystallization.

**A**

6FAM-βATTPNTG**GLFG**NTQT

6FAM-βATTPNTG**FSFG**NTQT

6FAM-βATTPNTGLS**FG**NTQT

**B**

*ISHM*WGSGTTPNTG**GLFG**NTQT$[GS]_{16}$K*GSGTTPNTG**GLFG**NTQT

*ISHM*WGSGTTPNTG**FSFG**NTQT$[GS]_{16}$K*GSGTTPNTG**FSFG**NTQT

*ISHM*WGSGTTPNTGLS**FG**NTQT$[GS]_{16}$K*GSGTTPNTGLS**FG**NTQT

**C**

*GSHM*WGC*GTTPNTG**GLFG**NTQT$[GS]_{16}$KGSGTTPNTG**GLFG**NTQT

**D**

DSG**GLFG**SK

DSG**FSFG**SK

DSGLS**FG**SK

**Figure S3. Affinities of GLFG-derived peptides to CA hexamers in the presence of *E. coli* lysate measured by MST**

Binding profiles of GLFG-peptides with single- (dotted line) and double-motifs (solid line) are shown in left. Apparent $K_D$ value ($K_{D,app}$) is presented (right). N.D. not determined. Data are expressed as mean ± s.e.

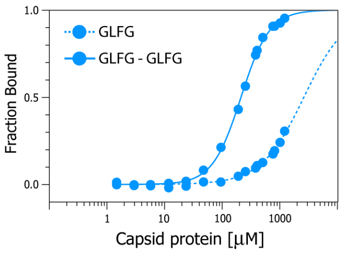


| | $K_{D,app}$ (mM) |
|---|---|
| GLFG | >2 |
| GLFG - GLFG | 0.212 ± 0.003 |

**Figure S4. Structures of CA hexamers with various FG peptides**
Overview of representative FG-binding sites (cartoon) with different FG peptides, GLFG, FSFG and FG (in stick representation). Six datasets (i-vi) of peptides at various apparent concentrations are indicated. (A) FG peptides superimposed on the Fo-Fc annealed omit map (orange, contoured at 2.0 and 1.5 σ for i-iv, vi and v respectively). (B) FG peptides superimposed on the 2Fo-Fc map (blue, contoured at 1.0 and 0.7 σ for i-iv, vi and v respectively). For clarity, electron densities for peptides are shown only.

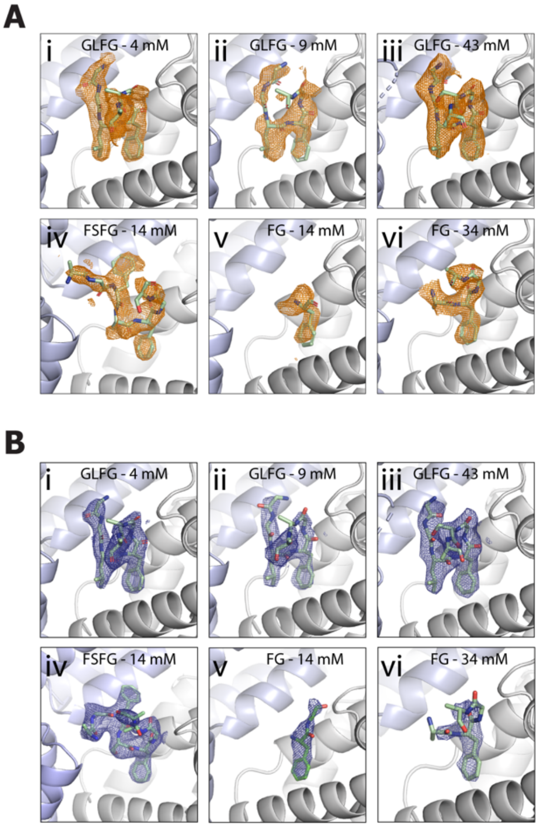

**Figure S5. Malleability of CA FG-binding pocket**
FG-binding pocket of CA in transparent surface representation. The flexibility of the Lys70 side chain (in white ball-and-stick; indicated) allows the pocket to remodel to accommodate the binding of GLFG- (A), FSFG- (B) and FG-derived peptides (C**)**. The peptides are shown in red and side chains for GLFG, FSFG and FG are shown in stick representation.

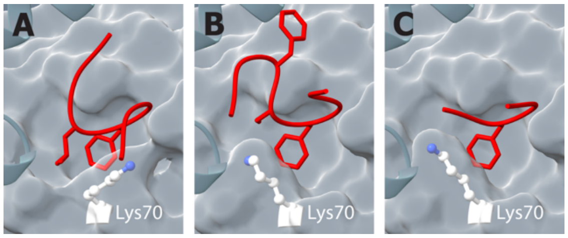

**Figure S6. “Footprints” of GLFG, FSFG, and FG motifs within the CA binding pocket**
Binding surface (red) of GLFG (A), FSFG (B) and FG (C) (in stick representation). Two adjacent protomers are in white and cyan respectively.

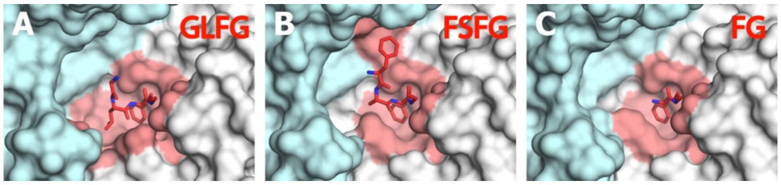

**Figure S7. The quality of core-like particles (CLPs) used for MST**

The micrograph of negatively stained CLPs obtained by transmission electron microscopy. Bar 500 nm.

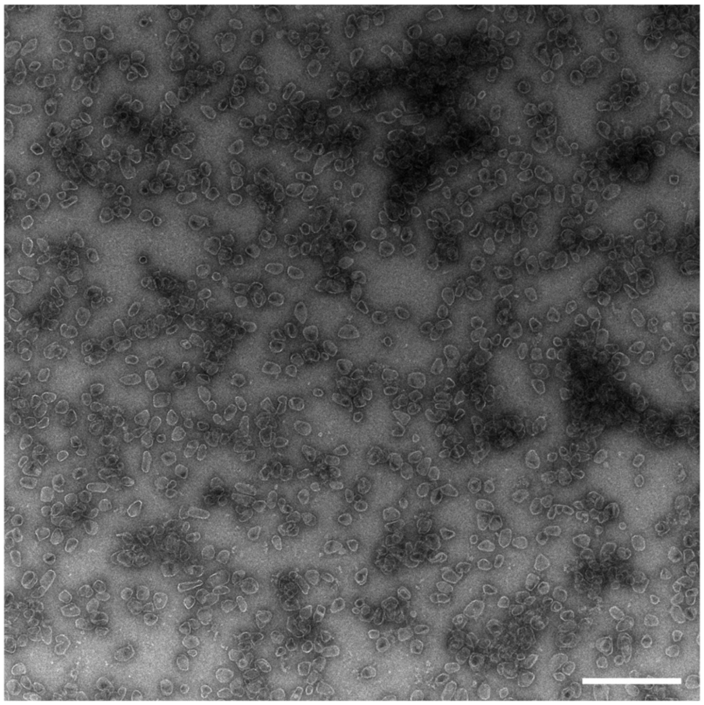

**Figure S8. Sequence analysis and design of FG-peptides with binding enhancers**
(A) Primary sequences of C-terminal regions of human NUP58, POM121 and NUP153. Basic residues are highlighted in red and FG motifs are indicated (black dots). Sequences underlaid with boxes represent those in engineered peptides (FTFG/FG in gray and basic-residues region in blue). (B) 6-Carboxyfluorescein (6-FAM) conjugated peptides used for MST assay and their designation. NUP58-FG•Cterm: $NUP58_{571\text{-}587/588\text{-}599}$, NUP58-FG: $NUP58_{571\text{-}587}$, NUP58-Cterm: $NUP58_{586\text{-}599}$, POM121-FG•Cterm: $POM121_{1212\text{-}1223/1236\text{-}1249}$, POM121-FG: $POM121_{1212\text{-}1223}$, POM121-Cterm: $POM121_{1234\text{-}1249}$, NUP153-FTFG•Cterm: $NUP153_{1411\text{-}1424/1465\text{-}1475}$, NUP153-FTFG: $NUP153_{1411\text{-}1424}$, and NUP153-Cterm: $NUP153_{1463\text{-}1475}$.

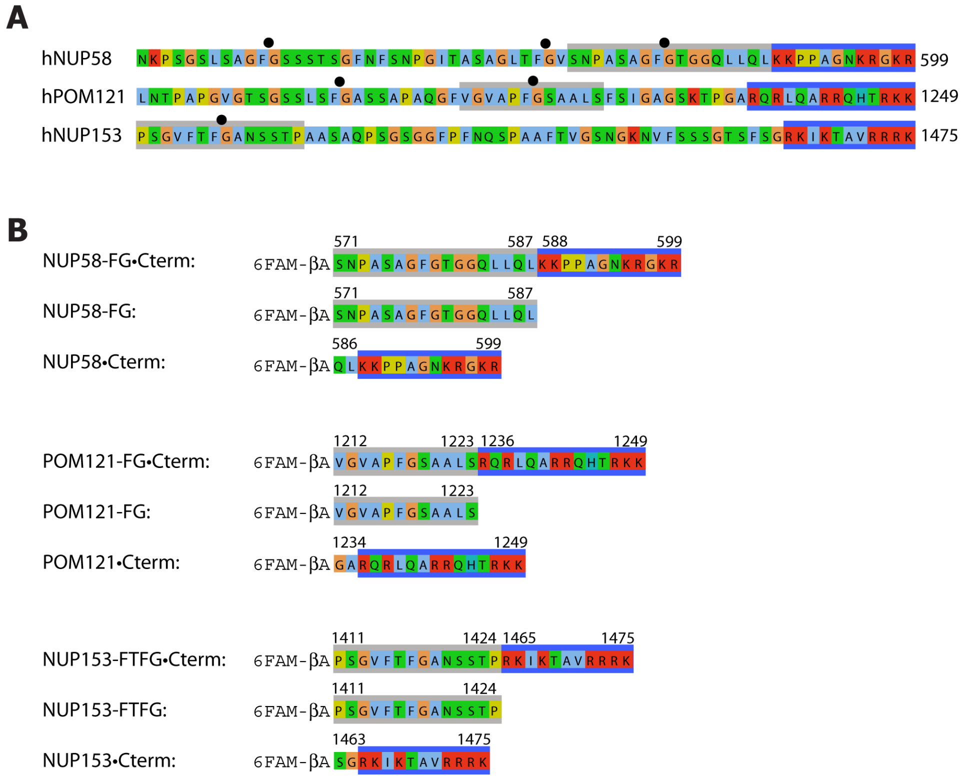

**Table S1. Summary of X-ray data collection and refinement statistics**

| | CA-GLFG$_{4\ mM}$ | CA-GLFG$_{9\ mM}$ | CA-GLFG$_{43\ mM}$ |
|---|---|---|---|
| ***Data collection*** | | | |
| X-ray source | BNL 17-ID-1 | BNL 17-ID-2 | BNL 17-ID-2 |
| Software | XDS | XDS | XDS |
| Space group | $P2_12_12_1$ | $P2_12_12_1$ | $P2_12_12_1$ |
| Unit cell dimensions | | | |
| *a*, *b*, *c* (Å) | 136.5 136.8 208.2 | 135.2 135.3 206.1 | 135.9 136.3 206.8 |
| α, β, γ (°) | 90.0 90.0 90.0 | 90.0 90.0 90.0 | 90.0 90.0 90.0 |
| ASU content | 2 | 2 | 2 |
| Wavelength (Å) | 0.920119 | 0.979338 | 0.920105 |
| Resolution range (Å)[a] | 34.70–3.00 (3.18–3.00) | 33.90–3.10 (3.28–3.10) | 34.20–3.00 (3.18–3.00) |
| $R_{sym}$ | 0.146 (0.838) | 0.153 (0.703) | 0.132 (0.769) |
| $R_{meas}$ | 0.163 (0.937) | 0.175 (0.817) | 0.151 (0.879) |
| $\langle I/\sigma I \rangle$ | 8.51 (1.91) | 7.46 (1.75) | 8.45 (1.75) |
| $CC_{1/2}$ (%) | 99.5 (61.4) | 99.3 (65.0) | 99.5 (64.9) |
| Completeness (%) | 98.5 (98.6) | 95.4 (96.2) | 99.5 (99.3) |
| Redundancy | 4.99 (5.02) | 3.96 (3.72) | 4.23 (4.31) |
| ***Refinement*** | | | |
| No. total reflections | 388,462 | 262,338 | 326,218 |
| No. unique reflections | 77,790 | 66,167 | 77,143 |
| No. test reflections[b] | 3,931 | 3,345 | 3,719 |
| $R_{work}$ / $R_{free}$ | 23.6 / 26.7 | 22.9 / 25.5 | 22.3 / 26.1 |
| No. atoms | 19,212 | 19,394 | 19,520 |
| Protein | 19,212 | 19,392 | 19,516 |
| Ion | 0 | 2 | 4 |
| Wilson B-factor (Å$^2$) | 66.1 | 63.2 | 68.4 |
| Average B-factors (Å$^2$) | 71.0 | 65.0 | 73.0 |
| RMS deviations | | | |
| Bond lengths (Å) | 0.002 | 0.002 | 0.002 |
| Bond angles (°) | 0.407 | 0.420 | 0.395 |
| ***MolProbity Statistics***[c] | | | |
| All atom clash score | 2.81 | 3.15 | 3.09 |
| Rotamer outliers (%) | 0.41 | 0.50 | 0.50 |
| Cβ deviations>0.25 Å | 0 | 0 | 0 |
| ***Ramachandran*** [c] | | | |
| Favored region (%) | 96.4 | 97.8 | 96.9 |
| Outliers (%) | 0 | 0 | 0 |
| ***PDB accession code*** | 9EDZ | 9EE0 | 9EE1 |

[a] Values in parentheses are for highest-resolution shell
[b] Random selection
[c] Values obtained from MOLPROBITY

**Table S1. Summary of X-ray data collection and refinement statistics**

| | $CA\text{-}FSFG_{14\ mM}$ | $CA\text{-}FG_{14\ mM}$ | $CA\text{-}FG_{34\ mM}$ |
|---|---|---|---|
| ***Data collection*** | | | |
| X-ray source | BNL 17-ID-1 | BNL 17-ID-1 | BNL 17-ID-1 |
| Software | XDS | XDS | XDS |
| Space group | $P2_12_12_1$ | $P2_12_12_1$ | $P2_12_12_1$ |
| Unit cell dimensions | | | |
| *a*, *b*, *c* (Å) | 136.4 136.4 208.8 | 135.6 135.9 206.5 | 135.9 136.4 207.3 |
| α, β, γ (°) | 90.0 90.0 90.0 | 90.0 90.0 90.0 | 90.0 90.0 90.0 |
| ASU content | 2 | 2 | 2 |
| Wavelength (Å) | 0.920119 | 0.920099 | 0.920099 |
| Resolution range (Å)[a] | 34.47–2.99 (3.17–2.99) | 34.01–3.10 (3.29–3.10) | 33.07–3.10 (3.29–3.10) |
| $R_{sym}$ | 0.124 ((>1) | 0.144 (0.495) | 0.148 (0.741) |
| $R_{meas}$ | 0.137 ((>1) | 0.170 (0.591) | 0.177 (0.894) |
| <I/σI> | 9.64 (1.59) | 6.44 (2.12) | 6.50 (1.81) |
| $CC_{1/2}$ (%) | 99.7(50.6) | 99.0 (73.0) | 99.1 (59.1) |
| Completeness (%) | 98.8 (97.7) | 96.0 (97.5) | 95.9 (96.8) |
| Redundancy | 5.71 (5.79) | 3.41 (3.30) | 2.92 (2.83) |
| ***Refinement*** | | | |
| No. total reflections | 446,517 | 228,320 | 197,493 |
| No. unique reflections | 78,145 | 67,035 | 67,605 |
| No. test reflections[b] | 3,959 | 3,391 | 3,440 |
| $R_{work}$ / $R_{free}$ | 23.3 / 25.4 | 22.8 / 26.1 | 21.9 / 25.2 |
| No. atoms | 19,425 | 19,261 | 19,347 |
| Protein | 19,424 | 19,260 | 19,345 |
| Ion | 1 | 1 | 2 |
| Wilson B-factor ($Å^2$) | 80.2 | 58.7 | 65.5 |
| Average B-factors ($Å^2$) | 90.0 | 60.0 | 68.0 |
| RMS deviations | | | |
| Bond lengths (Å) | 0.002 | 0.002 | 0.002 |
| Bond angles (°) | 0.391 | 0.405 | 0.405 |
| ***MolProbity Statistics***[c] | | | |
| All atom clash score | 2.28 | 2.43 | 2.76 |
| Rotamer outliers (%) | 0.54 | 0.30 | 0.74 |
| Cβ deviations>0.25 Å | 0 | 0 | 0 |
| ***Ramachandran***[c] | | | |
| Favored region (%) | 97.7 | 97.0 | 96.9 |
| Outliers (%) | 0 | 0 | 0 |
| ***PDB accession code*** | 9EE2 | 9EE3 | 9EE4 |

[a] Values in parentheses are for highest-resolution shell
[b] Random selection
[c] Values obtained from MOLPROBITY
